# PAG orchestrates T cell immune synapse function by binding to actin

**DOI:** 10.1101/2024.11.29.625440

**Authors:** Emily K. Moore, Marianne Strazza, Xizi Hu, Carly Tymm, Matthieu Paiola, Michael J. Shannon, Xinping Xie, Shoiab Bukhari, Shalom Lerrer, Emily M. Mace, Robert Winchester, Adam Mor

**Affiliations:** Columbia Center for Translational Immunology, Department of Medicine, Columbia University Medical Center; New York, NY 10032, USA; Division of Rheumatology, Department of Medicine, Columbia University Medical Center; New York, NY 10032, USA; Department of Pediatrics, Columbia University Medical Center, New York, NY 10032, USA; Laboratory of Synthetic Embryology, The Rockefeller University, New York, NY 10065, USA; Barnard College, Columbia University; New York, NY 10027, USA; Lymphoma Immunotherapy Program, Tisch Cancer Institute, Icahn School of Medicine at Mount Siani, New York, NY 10029, USA; Herbert Irving Comprehensive Cancer Center, Columbia University Medical Center; New York, NY 10032, USA

**Author notes:** Corresponding author: Adam Mor MD PhD.

## Abstract

Many immunotherapies impact T cell function by impacting the immune synapse. While immunotherapy is extremely successful in some patients, in many others, it fails to help or causes complications, including immune-related adverse events. Phosphoprotein Associated with Glycosphingolipid Rich Microdomains 1 (PAG) is a transmembrane scaffold protein with importance in T cell signaling. PAG has 10 tyrosine phosphorylation sites where many kinases and phosphatases bind. PAG is palmitoylated, so it localizes in lipid rafts of the membrane, and contains a C-terminal PDZ domain to link to the actin cytoskeleton. As a link between signaling-protein-rich membrane regions and the actin cytoskeleton, PAG is an exciting and novel target for manipulating immune function. Here, we sought to determine if PAG works with actin to control T cell synapse organization and function. We found that PAG and actin dynamics are tightly coordinated during synapse maturation. A PDZ domain mutation disrupts the PAG-actin interaction, significantly impairing synapse formation, stability, and function. To assess the impact of the PDZ mutation functionally in vivo, we employed a mouse model of type IV hypersensitivity and an OVA-tumor mouse model. In both systems, mice with T cells expressing PDZ-mutant PAG had diminished immune responses, including impaired cytotoxic function. These findings highlight the importance of the PAG-actin link for effective T cell immune synapse formation and function. The results of our study suggest that targeting PAG is a promising approach for modulating immune responses and treating immune-related diseases.

**One Sentence Summary:** Adaptor protein PAG links to the actin cytoskeleton, and this link is essential for T cell synapse formation and cytotoxic function.

## INTRODUCTION

To exert many of their effector functions, T cells must form healthy, stable immune synapses with antigen-presenting cells and target cells. The immune synapse is a complex and carefully organized structure that forms at the interface of two cells when T cell receptors (TCRs) encounter cognate MHC+peptides (*1*). The synapse is a dynamic structure. While actin continually polymerizes, some proteins migrate toward the center and others move toward the edges (*2–5*). Precise synapse organization is critical because proteins within a signaling pathway often require physical proximity (*6*). Thus, immune synapse organization has been a recent therapeutic target for cancer, autoimmune, or transplant patients (*7*).

Treatments targeting synapse proteins provide direct and nuanced manipulation of immune responses with promising results. The field of cancer immunotherapy has immensely improved outcomes for many cancer patients. One notable target for cancer is PD-1, an inhibitory receptor on antigen-activated T cells that is critical in the induction and maintenance of immune tolerance to self (*8*). Since tumor cells may overexpress PD-L1 to inhibit T cells through the PD-1 pathway by blocking the interactions of PD-1 and PD-L1, T cells remain actively able to target malignant cells (*9, 10*). However, overall, less than 20% of all cancer patients are estimated to respond to PD-1/PD-L1 inhibitors (*11*). One common complication in anti-PD-1 therapy is relapse. This may be because PD-1 signaling is also responsible for maintaining progenitor T cells (*12*). By blocking PD-1, we enhance the short-term immune response at the expense of the long-term immune reserve. Another common complication, especially among those whose cancers respond well, is immune-related adverse events: autoimmune symptoms following treatment with immune checkpoint inhibitors (*13, 14*). With further understanding of the complexity of the PD-1 pathway and the individual players that mediate different downstream effects, we will be better able to develop therapeutic approaches that enhance T cell anti-tumor responses without undesired outcomes, like progenitor depletion-induced relapse or immune-related adverse events (*15*).

Phosphoprotein associated with glycosphingolipid-rich microdomains 1 (PAG) is phosphorylated downstream of PD-1 ligation. This phosphorylation is required for the full impact of PD-1 signaling on many T cell effector functions (*16*). PAG is an inhibitory transmembrane adaptor protein (TRAP) with a 16-amino acid extracellular domain and a 397-amino acid cytoplasmic tail (in humans) (*17*). PAG is highly expressed in lymphocytes and is the only TRAP family protein that is also expressed in other tissues outside of blood cells (*18*). PAG localizes to lipid-rich membrane regions in T cells thanks to constitutive palmitoylation, and it recruits cytosolic kinases and phosphatases to its 10 tyrosine phosphorylation sites to participate in outside-in signaling cascades (*17, 19*). For example, when CSK, an inhibitory tyrosine kinase, binds to phosphorylated PAG, it is brought in proximity to inactivate the tyrosine kinase LCK to prevent signaling through the TCR (*17, 19–22*). Other binding partners include the SRC family kinases SRC, FYN, and LYN, among other kinases and phosphatases (*23–25*). Thus, PAG is essential in controlling the clustering of synapse-related signaling molecules.

Of importance to this study, PAG also has a C-terminal PDZ domain that binds to EBP50, a protein that connects to the actin cytoskeleton via Ezrin (*19, 26, 27*). PDZ domain proteins interact with the cytoskeleton, and they are essential to recruit and organize membrane receptors in all cell types (*28*). PDZ-domain proteins were first studied in the context of neuronal synapses and epithelial cells, and more recent studies have begun to suggest likely importance in immune synapse formation and function (*29*). For example, Scrib and Dlg1 have been shown to concentrate at the point of initial contact of the T cell with an antigen presenting cell, but once the TCR concentrates in the immune synapse, Scrib and Dlg1 instead enrich at the opposite pole of the cell (*30*). Most PDZ domain containing proteins are cytoplasmic, including EBP50, Scrib, and Dlg1, and must associate with the cell membrane through variable post-translational modifications or interactions with other proteins (*31, 32*). Uniquely, PAG is the only PDZ-containing protein that is transmembrane.

Actin is a major cytoskeletal protein essential for many cellular functions, including immune synapse formation (*33*). Actin dynamics coordinate the movement of molecules within the synapse, for example, clustering TCRs toward the center to accumulate a strong enough signal to activate the T cell (*34*). Since actin is essential in synapse organization, we must understand how actin polymerization affects the localization of other proteins, and whether PAG may play a role.

Cholesterol-rich membrane regions, or lipid rafts, organize membrane proteins to facilitate signal transduction and amplification (*35*). Lipid rafts are known to be necessarily enriched around the TCR after its engagement (*36, 37*). We hypothesized that PAG is a critical understudied link between actin and other lipid-raft proteins essential for immune synapse signaling and dynamic organization. Because of the importance of subcellular localization of signaling complexes and the complexity of healthy synapse formation in T cell responses, the PAG-actin link may have many critical functional implications.

The studies here help us understand PAG’s role in coordinating T cell immune synapse organization as a link between plasma membrane lipid rafts and the actin cytoskeleton. PAG has shown promise as a target for therapeutic antibodies in mouse studies (*38*). The anti-PAG antibody results in mislocalization of PAG within the synapse. This study helps elucidate the impact that PAG mislocalization could have on T cell synapse organization and function. We show that the PAG-actin link is essential for T cell synapse formation and function, with macroscopic impacts on the immune response to haptens and tumors in vivo.

## RESULTS

### PAG is a dynamic member of the immune synapse

PAG is seen evenly distributed around the plasma membrane of a resting T cell. When the T cell forms an immune synapse with an antigen presenting cell or target cell, PAG is more highly concentrated within the immune synapse than the opposite pole of the cell (*16*). To more closely examine the localization of PAG as the immune synapse forms and matures, we performed live cell imaging of Jurkat T cells expressing PAG-GFP, co-cultured with SEE-loaded Raji B cells. When a T cell initiates the formation of an immune synapse, PAG is enriched at the point of initial cell-cell contact (Fig. 1A-B; upper). As the immune synapse progresses, the contact area between the T and target cells grows, and PAG is enriched throughout the contact area (Fig. 1A-B; middle). In a fully mature immune synapse, the T cell may be seen wrapping around its target (Fig. 1A; lower). In a mature synapse, PAG can be found at the highest concentration at the periphery of the synapse (Fig. 1B; lower & Fig. 1C). The dynamic localization of PAG in the immune synapse is akin to that of actin. As the synapse matures, actin rapidly clears almost entirely from the center of the synapse (*2*).

**Fig. 1.**
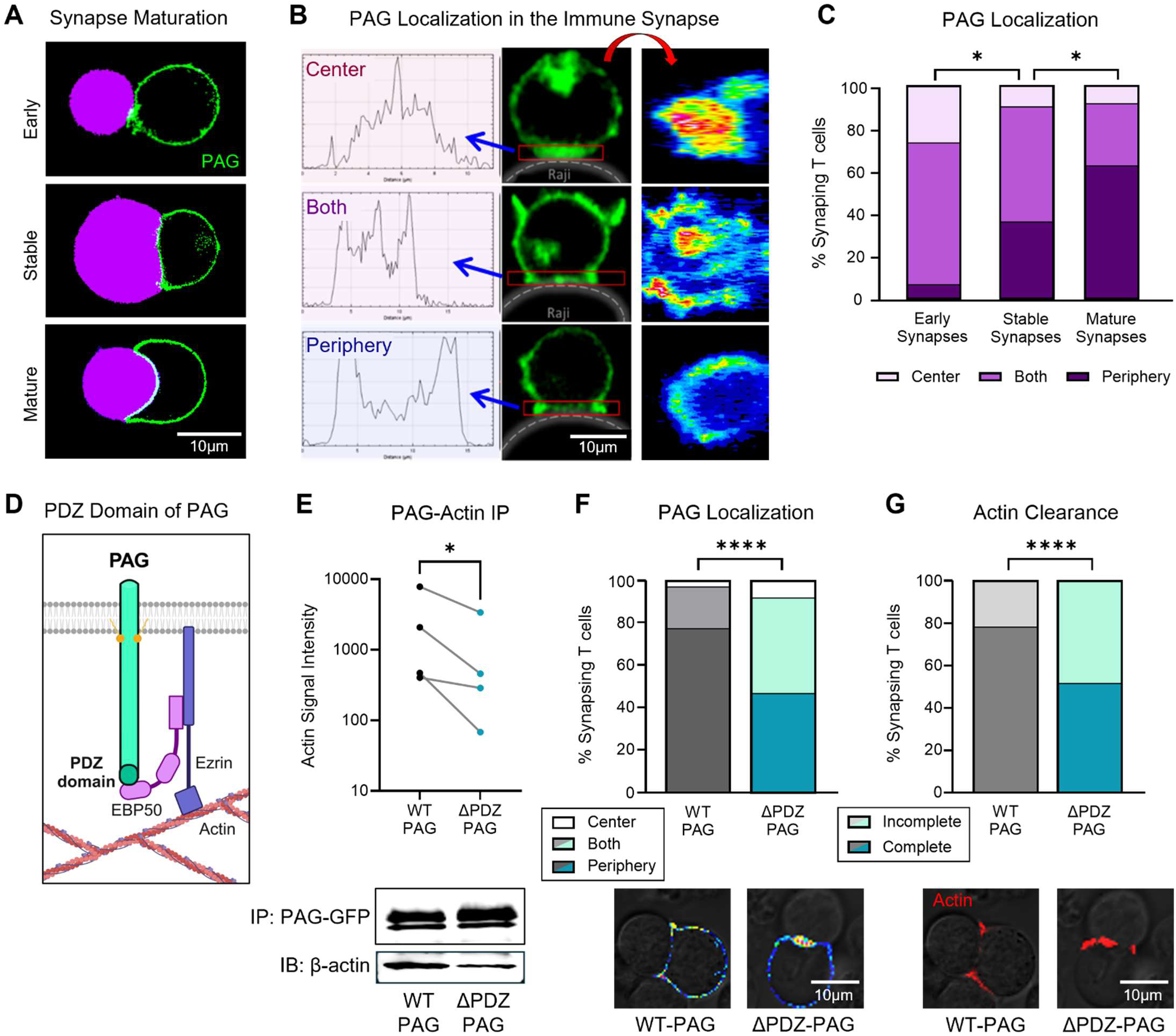
PAG is a dynamic member of the immune synapse, and the PDZ domain of PAG links to the actin cytoskeleton. **(A-C)** Jurkat T cells were transduced with PAG-GFP (green) and incubated with SEE-loaded Raji T cells (violet). Images were captured using ZEISS LSM 700, and image analysis was completed using the Plot Profile tool on ImageJ. **(A)** Representative images. 1.0cm = 10µm. **Upper**: Early immune synapse. **Middle**: Stable immune synapse. **Lower**: Mature immune synapse. **(B-C)** PAG localization was categorized based on the area of the synapse with the greatest intensity. **(B)** Representative images. 0.89cm = 10µm. **Upper**: Cells were categorized as “Center” if PAG intensity was greatest at the center of the immune synapse. **Middle**: Cells were categorized as “Both” if PAG intensity was greatest at the edges and center or throughout the full synapse interface. **Lower**: Cells were categorized as “Periphery” if the PAG intensity was greatest at the periphery of the synapse. **Left**: Line analysis of PAG localization in the immune synapse. **Middle**: PAG in the immune synapse shown in the original XY-plane. **Right**: PAG in the immune synapse reconstructed in the XZ-plane using the orthogonal view of the Z stack (pseudocolor, red = highest intensity). **(C)** Each T cell was classified as early, stable, or mature based on the morphology of the synapse, and PAG localization was categorized as center, periphery, or both based on the area of highest intensity. The percentage of cells with each PAG localization pattern is shown within each synapse category. Statistics by chi-square test to compare Early to Stable (chi-square = 8.353, df = 2) and Mature to Stable (chi-square = 7.463, df = 2). N = 7 independent experiments, a total of 100 cells were analyzed. **(D)** Diagram of the link between PAG and actin. The PDZ domain of PAG binds the PDZ domain of EBP50. EBP50 binds to Ezrin, and Ezrin binds actin. **(E)** Co-immunoprecipitation for PAG-GFP and actin with lysate from Jurkat cells with WT- or ΔPDZ-PAG. **Upper**: Statistics by ratio paired t-test. N = 4 independent experiments. **Lower**: Representative Western blot. **(F-G)** Assay as described in A-C to compare PAG and actin localization in synapsing Jurkat T cells expressing WT-PAG-GFP versus ΔPDZ-PAG-GFP. N = 4 independent experiments, a total of 230 cells were analyzed. **Upper**: Statistics by chi-square test. **Lower**: Representative images of T cells with WT-PAG-GFP (left) and ΔPDZ-PAG-GFP (right). 0.76cm = 10µm. **(F)** PAG localization, pseudocolor (red = highest intensity). Chi-square = 67.223, degrees of freedom (df) = 2. **(G)** Actin localization was visualized using LifeAct (red), and localization was categorized “Complete” if the actin signal was confined to the periphery of the synapse or “Incomplete” if actin fluorescence in the center of the synapse was greater than the average signal within the cell. Chi-square = 51.987, df = 1. ⇑ = P ≤ 0.05, *** = P ≤ 0.0001.

Given the similarities in the localization of PAG and actin in the immune synapse, we hypothesized that actin may be controlling PAG’s localization via its link to the PDZ domain of PAG. To test this, we mutated PAG in the PDZ domain to prevent its link to actin and assess the impact on the localization of these proteins in the immune synapse. The PDZ domain of PAG binds to a PDZ domain in EBP50, which in turn binds Ezrin to connect to the actin cytoskeleton (Fig. 1D). We introduced a single amino acid substitution in PAG at position 429 within the PDZ domain to replace isoleucine with alanine (I429A) (ΔPDZ-PAG). This mutation was selected because it has been shown to prevent PAG from binding to EBP50 (*26*). Jurkat T cells were engineered to knock out endogenous PAG using CRISPR. These cells were then rescued with either wildtype (WT) PAG-GFP or ΔPDZ-PAG-GFP using lentiviral transduction. The rescued cells were sorted to express approximately endogenous levels of PAG-GFP (Fig. S1A). We confirmed that ΔPDZ-PAG has a reduced association with actin by co-immunoprecipitation (Fig. 1E).

Using the Jurkat-Raji conjugate system (*39*), we assessed the impact of the ΔPDZ mutation on the localization of both PAG and actin. In T cells with WT-PAG, PAG is most enriched at the edges of a mature synapse in nearly 80% of cells. In contrast, in T cells with ΔPDZ-PAG, only half of the cells properly concentrate PAG at the edges of the synapse (Fig. 1F; left). In the remainder of cells, PAG is found throughout the synapse or concentrated at the center of the synapse (Fig. 1F; right). Actin localization was similarly dysregulated in these cells. In T cells with WT-PAG, actin clears entirely from the center of the synapse in nearly 80% of cells (Fig. 1G; left). However, when PAG was mutated in the PDZ domain, we were surprised to see less consistent actin clearance, with only half of the cells properly clearing actin from the center of a mature T cell synapse (Fig. 1G; right). Thus, the PAG-actin link is essential for the proper dynamics of PAG localization and contributes to actin localization, either directly or indirectly.

### The PAG-actin link facilitates immune synapse formation

Seeing that the PDZ mutation of PAG impacted the localization of not only PAG but also actin in the immune synapse, we sought to characterize further the immune synapses formed by T cells expressing WT-PAG or ΔPDZ-PAG. Because we saw mislocalized actin in synapsing cells with ΔPDZ-PAG, we hypothesized that mutant PAG impairs the strength and maturity of an immune synapse. We imaged T cells placed on an antibody-coated coverslip to observe the complete immune synapse interface in greater detail. This immune synapse model uses anti-CD3 and anti-CD28 antibodies to induce T cell polarization and spread onto the coverslip, forming an immune synapse (*40*). When WT Jurkat T cells are placed on antibody-coated coverslips for 5 or 40 minutes, the T cell spreads on the coverslip. The area of the interface increases over time (Fig. 2A-B). Even more strikingly, the cell’s perimeter on the plate increases as actin reorganizes at the contact site (Fig. 2A, 2C). In contrast, T cells with PDZ-mutant PAG do not significantly increase their contact area from 5 to 40 minutes (Fig. 2A-B). They increase perimeter over time, but significantly less than WT-PAG cells (Fig. 2A, 2C), suggesting that ΔPDZ-PAG impairs immune synapse reorganization over time.

**Fig. 2.**
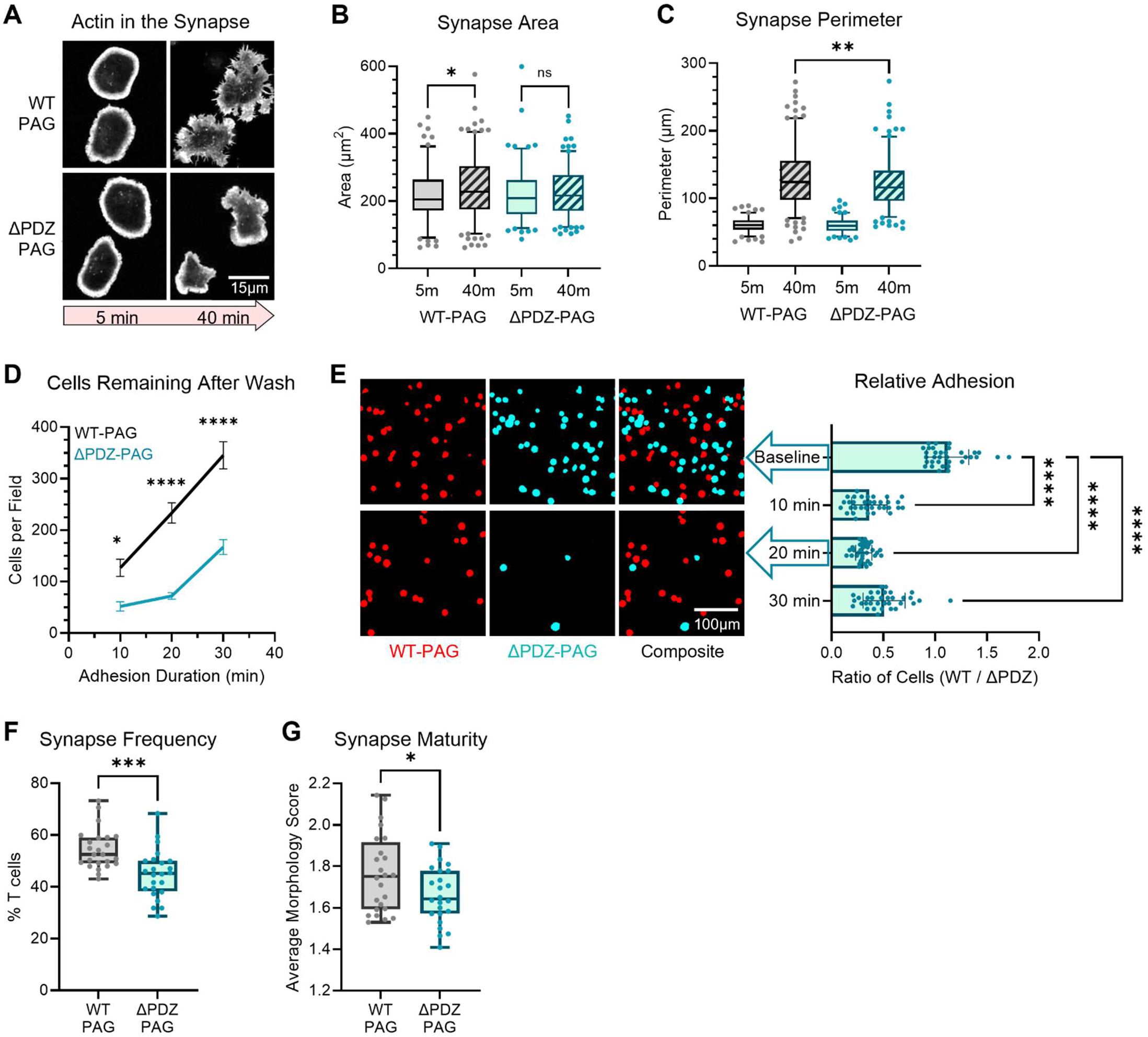
The PAG-actin link facilitates immune synapse formation. **(A-C)** Jurkat T cells with WT- or ΔPDZ-PAG-GFP were incubated for 5 or 40 min on anti-CD3 + anti-CD28 antibody-coated coverslips, fixed and stained for actin (phalloidin) and imaged with Zeiss LSM 900. N = 3 independent experiments. **(A)** Representative images showing actin localization. 0.66cm = 15µm. **(B)** Area of contact between the cell and the coverslip. **(C)** Perimeter of the contact area between the cell and the coverslip. 125-200 cells per condition. Statistics by ordinary one-way ANOVA with Fishers’ LSD test. **(D-E)** Pre-stimulated primary mouse CD8 T cells expressing WT- or ΔPDZ-PAG were differentially stained, mixed, and plated on mICAM1-coated coverslips. The plate was imaged, and after 10-30 min, the plate was washed once and imaged again. N = 2 independent experiments. **(D)** Cells per field were counted with Cell Profiler. Ordinary one-way ANOVA with Sidak’s multiple comparisons test. **(E) Left**: Representative images of WT and ΔPDZ-PAG T cells at baseline and after a 20-minute incubation + wash. 0.71cm = 100µm. **Right**: Ratio of cells per field. **(F-G)** Jurkat T cells with WT- or ΔPDZ-PAG-GFP were mixed with SEE-loaded Raji B cells and imaged live. N = 3 independent experiments. Statistics by unpaired t-test. **(F)** Percent of all T cells in a frame that are in an immune synapse at the imaging time, normalized to maintain an equal average WT percentage across experiments. **(G)** The morphology of each synapse was scored 1-3 for maturity, and scores were averaged across all cells in each field. **ns** = P > 0.05, ⇑ = P ≤ 0.05, ** = P ≤ 0.01, *** = P ≤ 0.001, *** = P ≤ 0.0001.

Since T cell spreading was impaired, we hypothesized that ΔPDZ-PAG may impair the strength of the immune synapse. To assess this, we used another synapse model to measure the strength of adhesion between T cells and their targets. The strength of an immune synapse can impact the length of time that the two cells remain in contact, as well as mechanical force-induced signaling. Together, this can mean the difference between weak TCR signaling not leading to T cell activation and robust TCR signaling, which results in an appropriate T cell response (*41–45*). To focus on the strength of LFA1-ICAM1 binding, which is essential for T cell synapse stability (*46, 47*), we coated a coverslip with murine ICAM1. We used primary mouse CD8 PAG-KO T cells rescued with WT-PAG or ΔPDZ-PAG (Fig. S1B), sorted by tdTomato expression, and preactivated with anti-CD3 and anti-CD28 antibodies. The two cell types were stained in different colors, mixed, and plated on the ICAM-coated coverslip. The cells were allowed to incubate for 10-30 minutes before washing the coverslip to remove any cells not strongly adherent to ICAM1. The longer the cells were incubated before washing, the more cells withstood washing, particularly for cells with WT-PAG (Fig. 2D). While the cells were plated at a 1:1 ratio, after one wash, more WT-PAG cells remained than ΔPDZ-PAG cells (Fig. 2E). This means that ΔPDZ-PAG T cells have weaker adherence to ICAM1. This suggests that linked PAG-actin dynamics facilitate formation of a strong and stable synapse.

Finally, we tested our hypothesis that ΔPDZ-PAG T cells would have impaired synapse generation by measuring how often a T cell is found in an immune synapse with a target cell and by assessing the morphology of those synapses. Jurkat T cells expressing WT- or ΔPDZ-PAG were mixed with SEE-loaded Raji target cells and imaged (*39*). The number of T cells in an immune synapse per field was compared to the total number of T cells in that field and normalized to the number of Raji cells in that field to find the percent of T cells in a synapse. We found a lower frequency of synapses among T cells expressing ΔPDZ-PAG compared to those with WT-PAG (Fig. 2F). When assessing the morphology of those synapses, we found that, on average, the synapses formed by ΔPDZ-PAG T cells were less mature than those formed by WT-PAG T cells (Fig. 2G). Based on these findings, we hypothesized that impaired synapse formation in ΔPDZ-PAG T cells would impede synapse-centric T cell functions, including activation state, ability to activate other immune cells, and cytotoxicity.

### The PAG-actin link in T cells contributes to tissue inflammation in contact dermatitis

We used a murine model of type IV hypersensitivity, or delayed hypersensitivity, to assess the function of ΔPDZ-PAG T cells in vivo. Type IV hypersensitivity, or delayed hypersensitivity, is a T cell-mediated form of hypersensitivity. In type IV hypersensitivity, the first exposure to an antigen results in T cell sensitization. Re-exposure can cause T cells to release cytokines to recruit other immune cells and induce inflammation (*48*). In humans, type IV hypersensitivity can result in allergic contact dermatitis (ACD).

Here, we induce ACD using the hapten 2,4-dinitrofluorobenzene (DNFB) (*49*). Because we saw impaired synapse formation in T cells with ΔPDZ-PAG, we hypothesized that the inflammatory response to DNFB would be lessened in mice with ΔPDZ-PAG T cells compared to mice with WT-PAG T cells. PAG-KO T cells were isolated from splenocytes and transduced to express WT-PAG or ΔPDZ-PAG (Fig. S1B). These cells were transferred to PAG-KO mice. The following day, the mice were sensitized with DNFB on their abdomen, and five days later, they were re-challenged with DNFB on their right ear. Two days after the challenge, the mice were assessed by histology and flow cytometry (Fig. 3A, Fig. S2A-C).

**Fig. 3.**
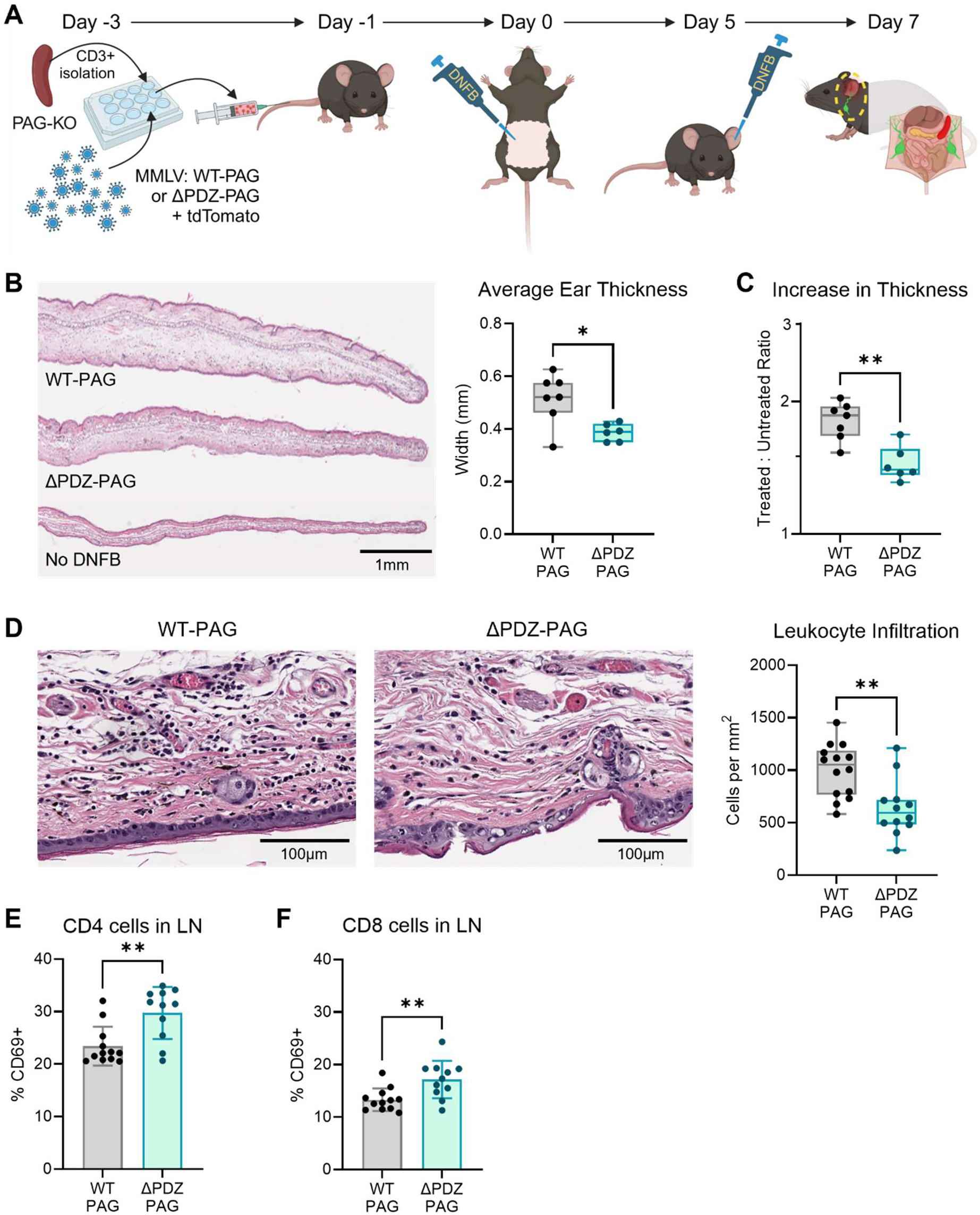
The PAG-actin link in T cells contributes to tissue inflammation in contact dermatitis. **(A)** Experiment design. PAG-KO CD3+ splenocytes were transduced with WT- or ΔPDZ-PAG + TdTomato and transferred into a WT mouse. The mouse was sensitized with DNFB on the abdomen and challenged with DNFB on the ear five days later. On day 7, ears were assessed by histology, and LNs were assessed by flow cytometry. N = 6-7 mice per group across 3 independent experiments. **(B)** Ear tissue was paraffin-embedded and stained with H&E. **Left**: Representative image. 1.1cm = 1mm. **Right**: Thickness was averaged from 9 equally spaced measurements along each section, with two sections per mouse. Statistics by Mann-Whitney t-test. **(C)** Ears were measured by micrometer three times on Day 7, and measurements were averaged. The thickness of the treated ear was normalized to the thickness of the contralateral untreated ear. Statistics by Mann-Whitney t-test. **(D)** Ear tissue was paraffin-embedded and stained with H&E. Dark round nuclei were counted and divided by the total area of tissue analyzed to find the count per mm^2^. Counts were normalized to maintain a consistent average WT value across experiments. 1.4cm = 100µm **(E-F)** Inguinal and cervical LNs were processed and stained for flow cytometry. Percent of CD69+ cells, gated on **(E)** CD3+CD4+ or **(F)** CD3+CD8+ live lymphocytes. Statistics by Mann-Whitney t-test. ⇑ = P ≤ 0.05, ** = P ≤ 0.01.

Compared to the contralateral untreated ear, mice with WT-PAG and mice with ΔPDZ-PAG all showed inflammation in response to re-challenge with DNFB. However, mice with WT-PAG T cells had significantly increased thickness of their treated ear compared to mice with ΔPDZ-PAG T cells (Fig. 3B-C, Fig. S2D). Upon greater magnification of H&E-stained ears, mice with WT-PAG T cells have a significantly higher density of infiltrating leukocytes (Fig. 3D). Flow cytometry was performed on T cells from the inguinal and cervical lymph nodes (LNs), which are the draining LNs for the sensitization site and the re-challenge site, respectively. Compared to WT-PAG mice, CD4 (Fig. 3E) and CD8 (Fig. 3F) T cells from ΔPDZ-PAG mice have a higher prevalence of the CD69 receptor, an activation marker. Thus, despite showing more activation markers in draining LNs, mice with ΔPDZ-PAG have an impaired inflammatory response to DNFB-induced hypersensitivity dermatitis, suggesting that despite LN T cell activation, ΔPDZ-PAG T cells have a diminished ability to mediate a type IV hypersensitivity response.

### The PDZ domain of PAG is required for CD8 T cell control of tumor growth but does not impact tumor infiltration

We used a mouse tumor model to assess further the impact of the PDZ mutation in PAG on the function of T cells in an immune response. A tumor model allows us to indirectly evaluate T cell killing of cancer cells, which requires a productive immune synapse. Additionally, it is valuable to re-assess PAG’s role in the T cell anti-tumor response since PAG is essential for PD-1 signaling (*16*) and has been targeted therapeutically in a mouse model (*38*). Here, we implanted a mouse colorectal carcinoma cell line that expresses the SIINFEKL peptide from ovalbumin protein (MC38-OVA) into WT CD45.1 mice. We then isolated CD8 T cells from PAG-KO OT-1 CD45.2 mice (donor cells) and transduced these cells to express tdTomato plus WT-PAG or ΔPDZ-PAG (Fig. S1B). Cells were sorted for tdTomato expression and transferred into the mice growing MC38-OVA tumors. Tumors were measured leading up to and following the transfer of donor cells, and at the endpoint, tumors were assessed by histology. In addition, the spleen, LNs, and tumor-infiltrating lymphocytes (TILs) were assessed by flow cytometry (Fig. 4A). This model allows us to evaluate the impact of the PDZ domain in PAG in T cells in an antigen-specific response.

**Fig. 4.**
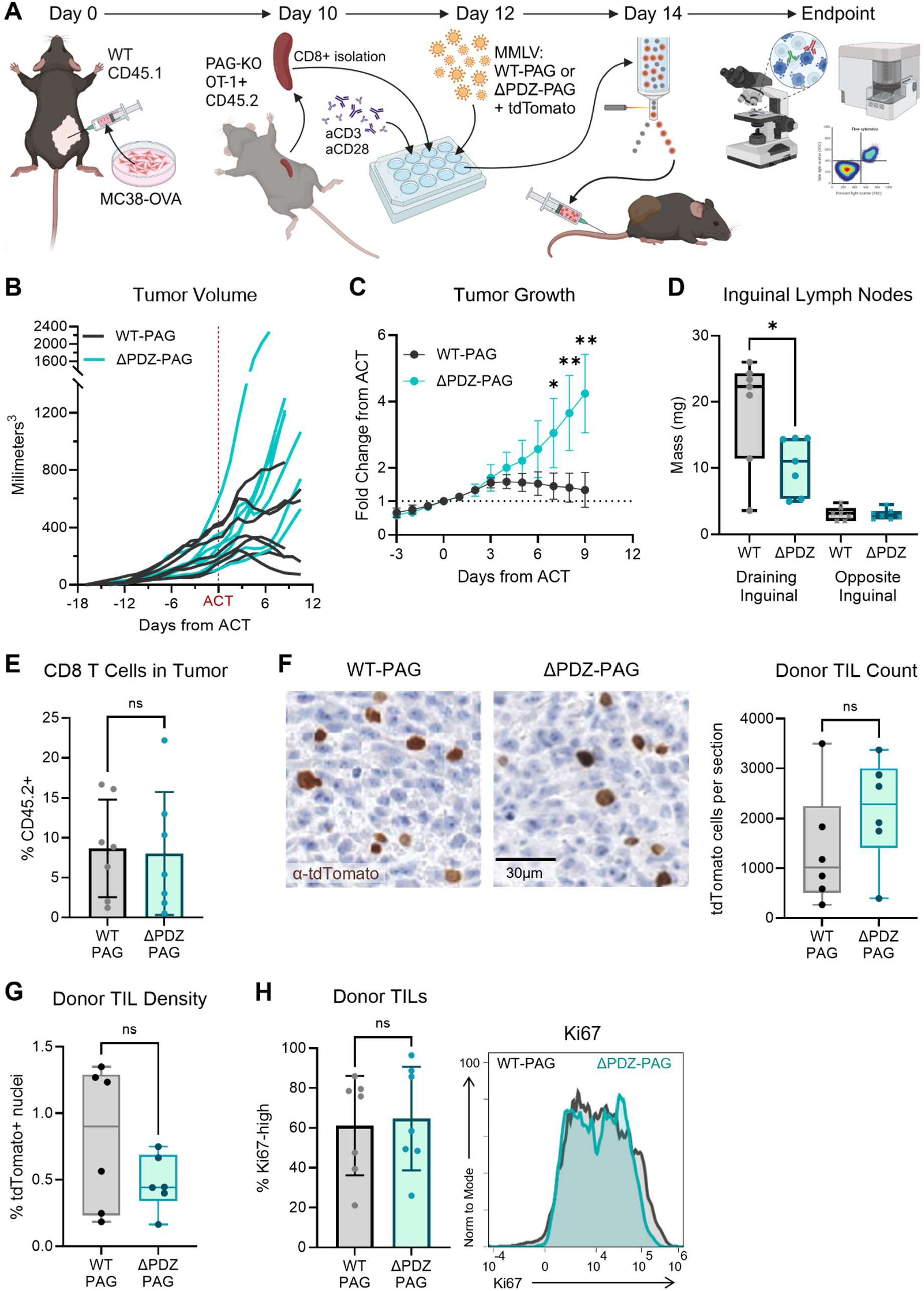
The PDZ domain of PAG is required for CD8 T cell control of tumor growth but does not impact tumor infiltration. **(A)** Experiment design. MC38-OVA tumor cells were implanted intradermally in WT CD45.1 mice. CD8+ splenocytes were isolated from PAG-KO CD45.2 mice, transduced with WT- or ΔPDZ-PAG + tdTomato, and sorted for tdTomato expression. Tumor growth was measured before and after the adoptive transfer of tdTomato cells, and tumor and LN were assessed by histology and flow cytometry at the endpoint. N = 7 mice per group across 3 independent experiments. **(B)** Tumor volume was calculated based on length and width measurements by caliper every 1-2 days before and after the adoptive transfer of donor cells. **(C)** Tumor size was normalized to the volume on the day of adoptive transfer to compare relative growth. Statistics by Mann-Whitney t-test with Holm-Sidak correction for multiple comparisons. **(D)** The draining inguinal LN and contralateral inguinal LN were isolated by dissection and weighed. Statistics by Welch’s unpaired t-test. (**E)** TILs were gated on CD8+ cells, and the percentage of those from the donor (CD45.2) is shown. Statistics by Mann-Whitney t-test. **(F-G)** Tumors were fixed and paraffin-embedded. Sections were stained with anti-tdTomato and hematoxylin. Statistics by Mann-Whitney t-test. (**F) Left**: Representative images. 1.0cm = 30µm. **Right**: The number of tdTomato+ cells per tumor section was counted using HALO. **(G)** The number of tdTomato+ cells was normalized to tumor section size using the number of hematoxylin-positive nuclei. **(H)** TILs were gated on CD8+ CD45.2+ (donor) cells. **Left**: Percent of donor TILs expressing high levels of Ki67. Statistics by Mann-Whitney t-test. **Right**: Ki67 fluorescence intensity histogram of pooled samples. **ns** = P > 0.05, ⇑ = P ≤ 0.05, ** = P ≤ 0.01.

Following the adoptive transfer of donor cells, the tumor growth slowed or reversed in mice who received WT-PAG OT-1 T cells. Conversely, tumors continued growing in mice who received ΔPDZ-PAG OT-1 T cells (Fig. 4B). In both groups, the tumors continued to grow for 2-3 days following transfer. By day 4, tumor growth diverged. The tumor growth curve flattened and then inverted for mice with WT-PAG OT-1 T cells, while it maintained continuous growth in mice with ΔPDZ-PAG OT-1 T cells (Fig. 4C). This means that even with antigen-specific T cells, when PAG was mutated in the PDZ domain, control of tumor growth was impaired. Additionally, although there was no difference in spleen mass (Fig. S3A), the tumor-draining LNs were significantly larger in mice with WT-PAG OT-1 T cells compared to mice with ΔPDZ-PAG T cells (Fig. 4D). This further supports the conclusion that there is an overall difference in the immune response when PAG is mutated in the PDZ domain.

We next sought to identify what T cell functions were impaired in ΔPDZ-PAG T cells to result in strikingly different control of tumor growth. We hypothesized that this mutation either impaired T cell ability to reach the tumor or impaired T cell activity and cytotoxicity once in the tumor. To evaluate tumor infiltration, we measured the percent of infiltrating CD8 T cells from the donor rather than the host by flow cytometry based on CD45 isoform. The percent of CD8 T cells from the donor varied in all tumors, with no overall difference between WT-PAG T cells and ΔPDZ-PAG T cells reaching the tumor (Fig. 4E). To validate these findings further, we measured infiltration by staining for tdTomato+ cells in the tumor histology sections. Again, there was no consistent difference in infiltration of OT-1 T cells with WT-PAG versus ΔPDZ-PAG. There was a comparable number of tdTomato+ cells per tumor section (Fig. 4F) and no difference in the percent of nuclei in the tumor that were tdTomato+ (Fig. 4G). Notably, the Ki67 expression was also comparable between the two groups, meaning that an earlier or later time point would likely still show no difference in infiltration between WT-PAG and ΔPDZ-PAG T cells (Fig. 4H). Thus, we concluded that the difference in tumor control between WT-PAG and ΔPDZ-PAG T cells was not due to differences in tumor access but rather impairment in the T cell anti-tumor response once the T cells reached the tumor microenvironment.

### CD8 T cells with PDZ-mutant PAG are activated and express high levels of PD-1

Next, we sought to test the hypothesis that tumor-infiltrating ΔPDZ-PAG T cells were ineffectively controlling tumor growth because they were less activated than WT-PAG T cells. To do this, we enriched lymphocytes from tumor tissue and analyzed them by flow cytometry. After gating on donor CD8 TILs, our panel looked at eight markers of activation, exhaustion, and function. Overall, the tumor infiltrating ΔPDZ-PAG T cells were noticeably different from WT-PAG T cells based on principal components analysis (PCA) (Fig. 5A). In particular, activation marker CD25 and the chemokine receptor CX3CR1 were more expressed on tumor-infiltrating T cells with ΔPDZ-PAG than those with WT-PAG (Fig. 5B-C). Thus, the tumor infiltrating ΔPDZ-PAG T cells appear activated even though they could not control tumor growth in vivo. The most noticeable difference between WT-PAG T cells and ΔPDZ-PAG T cells was the PD-1 expression level. Though all cells were PD-1 positive, T cells with ΔPDZ-PAG consistently had higher PD-1 expression than T cells with WT-PAG (Fig. 5D). To further assess the cell profile differences between WT-PAG and ΔPDZ-PAG in OT-1 T cells, we performed unsupervised cluster analysis and UMAP visualization on our flow cytometry results. We found that the largest cluster, Cluster 1, was more highly populated by ΔPDZ-PAG T cells than WT-PAG T cells (Fig. 5E-F). Further, this cluster has the highest PD-1 of any cluster (Fig. S3B), driven by the high PD-1 levels specifically found in ΔPDZ-PAG cells (Fig. 5G). However, interestingly, Cluster 1 is also the cluster with the lowest Granzyme B (Fig. S3B). This result is consistent with findings in the hypersensitivity model, where ΔPDZ-PAG T cells also mounted a weaker immune response despite the larger presence of activation markers.

**Fig. 5.**
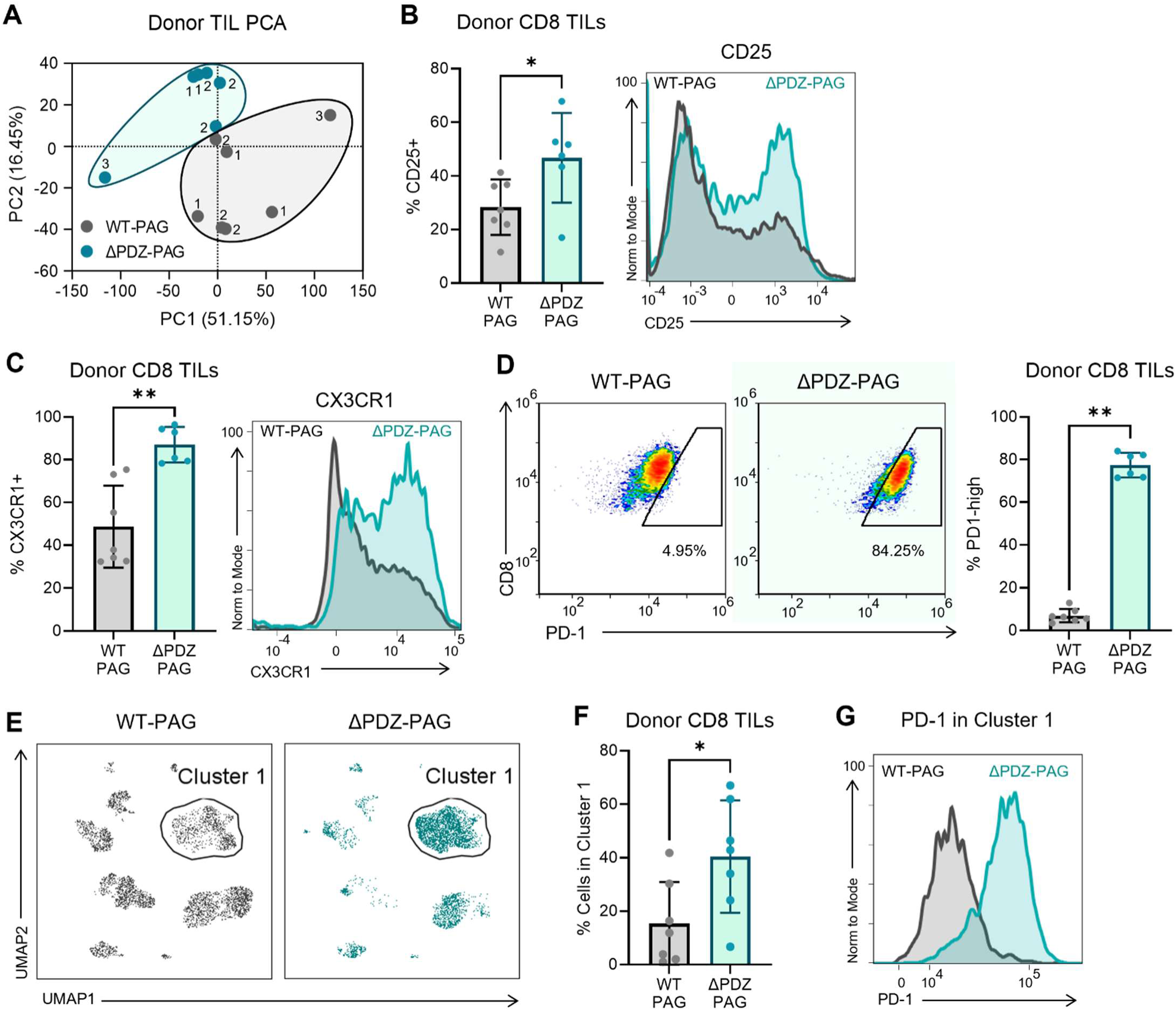
CD8 T cells with PDZ-mutant PAG are activated and express high levels of PD-1. N = 7 mice per group across 3 independent experiments. **(A)** Principal component analysis (PCA) of the normalized MFIs of 8 parameters of the flow cytometry results in donor TILs. **(B)** Donor TIL flow cytometry. **Left**: The percent expressing CD25. Statistics by Mann-Whitney t-test. **Right**: CD25 fluorescence intensity histogram of pooled samples. **(C)** Donor TIL flow cytometry. **Left**: The percent expressing CX3CR1. Statistics by Mann-Whitney t-test. **Right**: CX3CR1 fluorescence intensity histogram of pooled samples. **(D)** Percent of donor TILs expressing high levels of PD-1. **Left**: Representative flow cytometry plot of WT- and ΔPDZ-PAG TILs. **Right**: Statistics by Mann-Whitney t-test. **(E)** UMAP analysis with supervised clustering of 13 parameters and 9 clusters. **(F)** Percent of cells per sample found in cluster 1. Statistics by Mann-Whitney t-test. **(G)** PD-1 fluorescence intensity histogram for pooled sample cells in cluster 1. ⇑ = P ≤ 0.05, ** = P ≤ 0.01.

Notably, in our tumor model, the differences in activation marker expression are only seen among the tumor-infiltrating T cells. Donor T cells found in the draining LN showed contrasting trends. The draining LN from mice with ΔPDZ-PAG has a lower prevalence of donor OT-1 T cells (Fig. S3C). In contrast to what was seen in the tumor, CD25 and CX3CR1 are lower in ΔPDZ-PAG T cells than in WT-PAG T cells found in the draining LN (Fig. S3D-E). Additionally, when WT-PAG and ΔPDZ-PAG T cells are stimulated in vitro with anti-CD3 and anti-CD28 antibodies, there is no difference in their expression of these activation markers, including PD-1 (Fig. S3F). Because of this discrepancy, we suspect that ΔPDZ-PAG T cells interact with host immune and stromal cells differently than WT-PAG T cells, impacting their activation and function particularly in vivo.

### CD8 T cells with PDZ-mutant PAG have intact epithelial adhesion but have impaired cytotoxicity

To further understand why ΔPDZ-PAG cells could not control tumor growth, we performed several in vitro experiments to isolate and test specific effector functions. We saw approximately equal infiltration of donor T cells into tumors. To further confirm the ability of ΔPDZ-PAG cells to migrate in vitro, we tested adhesion to epithelial cells under shear stress (*50*). Rather than static TCR-induced activation, as tested in Fig. 2, this assay assesses the ability of T cells to bind to epithelial cells under the shear stress present in circulation. Thus, this assay serves as a model to determine the T cell’s ability to “arrest” movement and resist blood flow in the first step of extravasation (*50*). We plated equal numbers of WT-PAG and ΔPDZ-PAG T cells on CHO epithelial cells. CHO cells express ICAM1, and there is 100% sequence homology for the first 277 amino acids of ICAM1 between mouse and Chinese hamster (NCBI alignment of NP_034623.1 and NP_001233619.1), covering and expanding far beyond the binding site for LFA1 in the first Ig-like domain of ICAM1 (*51*). After the T cells settled onto the CHO cells, we applied a media flow at a speed equivalent to what a T cell would experience in a capillary. After 5 minutes of flow, a comparable number of WT and ΔPDZ-PAG T cells remained adherent to CHO epithelial cells (Fig. 6A). The ratio of WT-PAG T cells to ΔPDZ-PAG T cells remained centered around 1:1 (Fig. 6B). From this, we conclude that although adhesion in an immune synapse is impaired, adhesion in the context of extravasation is intact in ΔPDZ-PAG T cells.

**Fig. 6.**
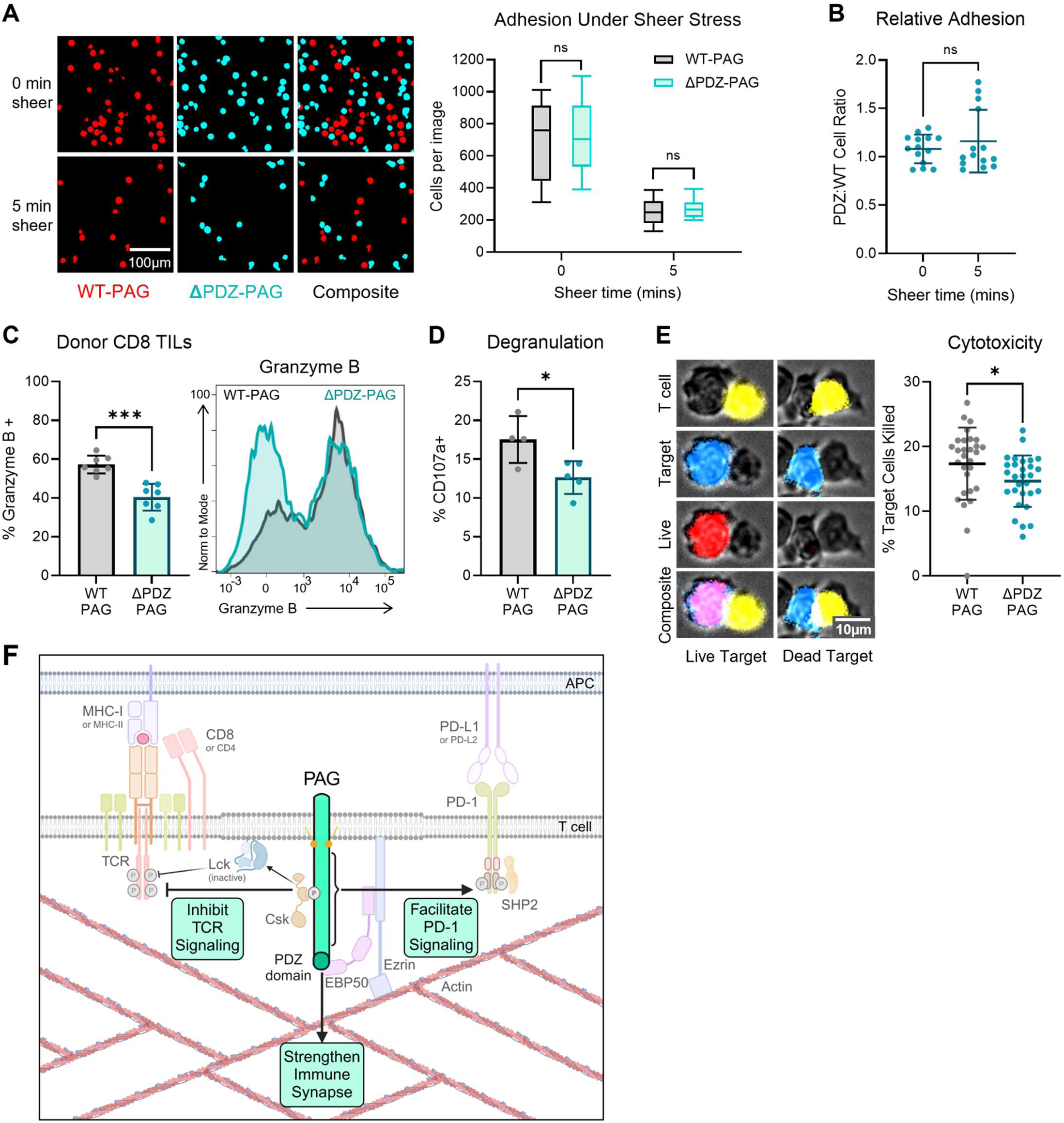
CD8 T cells with PDZ-mutant PAG have intact epithelial adhesion but have impaired cytotoxicity. **(A-B)** Murine CD8 T cells with WT-PAG and those with ΔPDZ-PAG were differentially stained, mixed, and plated on CHO cells and assessed for their resistance to shear stress. Statistics by Mann-Whitney t-test. N = 2 independent experiments. **(A) Right**: Representative images at baseline and after 5 min of media flow over the cells. 0.71cm = 100µm. **Left**: The number of each cell type was counted per 20x field. **(B)** The number of PDZ cells in each field was normalized to the number of WT cells in the corresponding field to assess the change in ratio with shear stress. **(C)** Donor TIL flow cytometry. **Left**: Percent expressing Granzyme B. Statistics by Mann-Whitney t-test. **Right**: PD-1 fluorescence intensity histogram of pooled samples. N = 7 mice per group across 3 independent experiments. **(D)** Percent of murine CD8 T cells with surface expression of degranulation marker CD107a following incubation with MC38-OVA target cells, measured by flow cytometry. Statistics by Welch’s unpaired t-test. N = 2 independent experiments. **(E)** Murine CD8 T cells with WT-PAG or ΔPDZ-PAG were mixed with Raji-SEE target cells and imaged after 24 hours of incubation. Live cells maintained staining with Calcein (Red), while apoptotic cells lost this dye. **Left**: Representative images. 0.61cm = 10µm. **Right**: The percentage of apoptotic target cells was normalized to maintain a consistent average for WT-PAG T cells between experiments. Statistics by unpaired t-test. N = 2 independent experiments. **(F)** PAG is a multifunctional protein that inhibits TCR signaling, facilitates PD-1 signaling, and strengthens the immune synapse. **ns** = P > 0.05, ⇑ = P ≤ 0.01, *** = P ≤ 0.001.

We next sought to assess the cytotoxic ability of ΔPDZ-PAG T cells compared to WT-PAG T cells. If the ΔPDZ-PAG T cells can get to the tumor and become activated, perhaps their impaired tumor control comes from a lack of ability to kill target cells. In the OVA tumor model, tumor-infiltrating donor T cells with ΔPDZ-PAG consistently have less granzyme B than WT-PAG T cells (Fig. 6C). This suggests that ΔPDZ-PAG T cells either have a lower cytotoxic capacity or degranulate more than WT-PAG T cells. We then looked at relative degranulation by measuring surface CD107a. Following culture with MC38-OVA tumor cells in vitro, ΔPDZ-PAG T cells have lower surface CD107a, suggesting less degranulation occurs in ΔPDZ-PAG T cells compared to WT-PAG T cells (Fig. 6D). Finally, we directly measured cytotoxicity by assessing target cell death after 24h culture with WT-PAG or ΔPDZ-PAG T cells. Compared to target cells cultured with WT-PAG T cells, fewer target cells died when cultured with ΔPDZ-PAG T cells (Fig 6E). These results suggest that mutation in the PDZ domain of PAG results in impaired cytotoxicity. Effective and directed cytotoxicity depends upon forming a productive immune synapse. Given the importance of PAG’s PDZ domain for strong immune synapses, T cells with a mutation in this domain are less able to kill antigen-specific tumor cells.

## DISCUSSION

In this study, we sought to identify the functional impact of PAG’s PDZ domain on synapse formation and T cell function. To do this, we mutated the PDZ domain in PAG and assessed the effect of this point mutation on T cell synapse formation and T cell function in vitro and in vivo. We showed in vitro that this mutation impacts the localization of both PAG and actin in an immune synapse and the T cell’s ability to form strong and mature immune synapses. In vivo, we showed that loss of the PDZ domain in PAG results in a diminished inflammatory response to DNFB-induced hypersensitivity. We also demonstrated that losing the PDZ domain in PAG prevents antigen-specific control of tumor growth in mice. Despite their inability to prevent tumor growth, we also noted that ΔPDZ-PAG T cells could migrate into tumor tissue. These TILs highly expressed activation markers, including higher levels of PD-1 than in WT-PAG T cells. An in vitro assessment of necessary T cell functions showed intact capacity to adhere to epithelial cells, which is the first step for extravasation and migration to the tumor. However, they showed decreased cytotoxic capability than WT-PAG T cells, which may help explain antigen-specific T cells’ lack of tumor control with ΔPDZ-PAG.

There are some limitations to our studies. It remains unclear the mechanism by which the PAG-actin link facilitates synapse formation. It could be that the actin network is directly distorted without the PAG link. However, the relationship could be indirect. Without the PAG link, PAG is mislocalized. Perhaps this results in abnormal TCR signaling, since PAG as an adaptor protein is not bringing important kinases and phosphatases where they need to be. Weaker TCR signaling and less cell activation, could cause incomplete activation of the Arp2/3 complex that nucleates actin polymerization and branching necessary for cell spreading during synapse maturation (*52–54*). Further, LFA1 may not efficiently switch to and maintain its high affinity state, impairing adhesion to ICAM1 (*55*). The relationship between actin and T cell signaling is complex and bidirectional.

Though cytotoxicity was mildly impaired in ΔPDZ-PAG T cells in vitro, we believe the functional impact of this mutation is more prominent in vivo. During an in vivo immune response, there are a series of interactions that T cells must make with other cells in their microenvironment (*56*). Since ΔPDZ-PAG impacts T cell synapse formation, there may be altered interactions with antigen presenting cells in the LNs, for example, which could result in weaker T cell targeting of tumor cells (*57*). Additionally, as the T cell interacts with its microenvironment, it also impacts the function of the cells around it. Weaker immune synapses with ΔPDZ-PAG T cells may directly or indirectly, through cytokine release, impair the activation of the cell on the other side of the synapse, thereby impacting the overall strength of the anti-tumor response (*58*).

Previous studies showed the necessity of PAG for PD-1 signaling. PAG knockdown cells lose PD-1-induced inhibition of IL-2, IFN-γ, adhesion, CD69 expression, and ERK phosphorylation (*16*). These functions were variably recovered when PAG KD cells were rescued with PAG mutated to prevent phosphorylation of different tyrosine sites (*16*). This suggests that these sites on PAG may dock various signaling mediators and that changed phosphorylation status in other conditions would lead to different effector functions. Further, patient data shows that higher expression of PAG is associated with worse outcomes in many cancers (*59–61*). In the murine MC38 colon adenocarcinoma tumor model, PAG KO mice have limited tumor growth and smaller tumors following anti-PD-1 treatment than WT mice (*16*). This correlated with increased CD3^+^ T cell infiltration into MC38 tumors in PAG KO mice and enhanced cytotoxicity of PAG KO murine T cells in vitro (*16*).

Due to its potential for coordinating signaling complexes around lipid-rich regions and its known role in the PD-1 pathway, PAG is an exciting and novel target for manipulating immune function. Prior therapeutic methods for cancer primarily focus on mimicking or disrupting ligand binding or enzyme function (*62–64*). PAG has no known ligands and is not known to have any intrinsic catalytic function (*17*). However, understanding PAG’s role in controlling T cell synapse organization opens more avenues for therapeutically targeting the immune synapse. Instead, targeting PAG with an antibody disrupts appropriate PAG localization (*38*). This is because a bulky antibody prevents PAG from locating within the narrow space of the synapse. Promisingly, mice treated with anti-PAG antibodies have slowed tumor growth (*38*). The results from the present study help illuminate how relocating PAG with an antibody will impact actin and which T cell functions will be impacted by PAG re-localization.

Based on this set of studies, we conclude that PAG is not simply an inhibitory protein, but a multi-functional protein. Intact PAG function is required for proper synapse formation. However, PAG’s inhibitory capacity can be measured upon effective synapse formation. Thus, PAG works as a modulator of T cell activation, facilitating T cell activation through synapse formation but preventing unnecessary or excessive TCR signaling and cell activation.

## MATERIALS AND METHODS

### Cell culture and mice

Jurkat T cells (ATCC) were cultured in RPMI medium (Corning 10-040-CM) supplemented with 10% FBS (Gibco 10438-026) and 1% Pen/Strep (P/S, 10,000 U/ml stock) (Corning 30-002-CI). Primary mouse T cells were isolated from spleen and inguinal LN of C57BL/6 (B6) PAG knockout (KO) mice or OT-1-B6 PAG KO mice (Alexander Tarakhovsky, Rockefeller Institute). Spleens and LN cells were dissociated through a 70-micron filter (Genesee Scientific 25-376) and rinsed with PBS (Corning 21-040-CV). Cells were washed (Eppendorf 5810R) and resuspended in MojoSort buffer (PBS with 0.5% BSA and 2mM EDTA), and CD3 or CD8 cells were isolated negative selection following the protocol of MojoSort kits (BioLegend 480024; BioLengend 480008). Isolated mouse T cells were cultured in T cell media: RPMI, 10% FBS, 1% P/S, 1x BME (MP Biomedicals 219024280), 1x GlutaMAX (Gibco 35050061), 1x Sodium Pyruvate (Gibco 11360070), 1x NEAA (Gibco 11140050), 100 IU/mL hIL2 (Gibco 200-02).

### CRISPR PAG KO in Jurkat cells

PAG KO Jurkat T cells were generated using CRISPR-Cas9 (*65*). Briefly, guide RNA (gRNA) was assembled by mixing equal amounts of 200µM PAG1-targeting CRISPR RNA (crRNA) (IDT) and 200µM trans-activating CRISPR RNA (tracrRNA, IDT 1072532). crRNA sequences were Hs.Cas9.PAG1.1.AA: 5’-CACGGCGAGAAGUGUGGACGGUUUUAGA GCUAUGCU-3’ and Hs.Cas9.PAG1.1.AB: 5’-UGGUCCCCACUAUGCUGUCGGUUUUA GAGCUAUGCU-3’.

gRNA was incubated for 5m at 95C and cooled to room temperature. An equal volume of 20µM Cas9 enzyme (IDT 1081058) was mixed with the gRNA and incubated at 37C for 15 min to form ribonucleoprotein. 3µL of gRNA was mixed with 1µL electroporation enhancer (IDT 1075915) and 20µL of Jurkat T cells (1 x 10^5^) in nucleofection solution (Lonza V4XC-1032). The mixture was transferred to a 16-well Nucleocuvette Strip and electroporated with the 4D-Nucleofector (Lonza) set to CL-120. Immediately following electroporation, 150µL of warm RPMI-10 was added, and cells were transferred to a 96-well plate for incubation at 37C.

After 48h, KO efficacy was assessed by Tracking of Indels by Decomposition (TIDE) analysis (*66*). Genomic DNA was isolated following protocol from the DNeasy mini kit (QIAGEN 69504). PCR was performed following the protocol for the Phusion Flash High-Fidelity PCR Master mix (Thermo Scientific F548S) with primers designed to flank the editing site (IDT FwdAA: 5’-GTGTGTGCTGCTTTAGTTCTTTC-3’, RevAA: 5’-CAGTGCCCACCTTGTTAGTT-3’, FwdAB: 5’-GAAACACAGAGGTCACCATAGG-3’, RevAB: 5’-GCAAACGGGCCATCAAATC-3’). The PCR product was converted to heteroduplexes and sequenced (Azenta/GENEWIZ). Sequencing results were analyzed using TIDE. The commercially available PAG antibody binds intracellularly so cells could not be sorted based on PAG level. Jurkat T cells do not survive well when plated as single cells, so the CRISPR cells were diluted to a few cells per well to form polyclonal colonies. After the pools grew, they were analyzed by flow cytometry, and the best ones were sequenced again and analyzed by TIDE and ICE analysis (*67*) to select one with the lowest PAG level. These cells were frozen into many tubes, and a new stock vial was thawed after 3-4 weeks of culture to minimize drift in PAG expression over time.

### Expressing WT-PAG-PAG and ΔPDZ-PAG-GFP in Jurkat cells

Transient transfections were done with the pEGFP-N1-PAG construct (*16*) and its derivates. pEGFP-N1-ΔPDZ-PAG-GFP was generated by mutating amino acid 429 from isoleucine to alanine by site-directed mutagenesis. Base pair 1285 was changed from adenine to guanine, and base pair 1286 was changed from thymine to cytosine. Site-directed mutagenesis was done following the protocol and reagents from QuikChange Lightning Site-Directed Mutagenesis Kit (Agilent 210519), and the following primers (IDT) were used: 5’-AGTAAACCTGAAGCTGCTGGGAGTAATTCCCATATAC-3’ and 5’-GTATATGGGAATTACTCCCAGCAGCTTCAGGTTTACT-3’. The sequence of the products was verified by Sanger sequencing with primer 5’-GAAAACCTTCAGGAGAAGGAAGGG-3’ (IDT).

Transient transfection of Jurkat cells was accomplished by electroporation using the AMAXA setting S018 on Nucleofector 2b (LONZA AAB-1001) with the reagents and protocol from the Cell Line Nucleofector Kit V (LONZA VCA-1003) or Ingenio Electroporation Kit for AMAXA (Mirus, MIR 50118).

The stable expression of WT-PAG or ΔPDZ-PAG in Jurkat T cells was accomplished using lentiviral transduction. WT-PAG-GFP and ΔPDZ-PAG-GFP were cloned into the pHR vector backbone using restriction cloning with EcoRI (Thermo Scientific FD0274), NotI (Thermo Scientific FD0593), and the Thermo FastDigest reagents and protocol. Note that the pHR vector had a second EcoRI site removed using site-directed mutagenesis. Products were sequence verified by Sanger sequencing (Azenta/GENEWIZ). Lentivirus was produced by transfecting 293T cells with 60µL of SuperFect Transfection reagent/protocol and 1.75µg pMD2G (Marks Philips, NYU), 3.25µg psPAX2 (Mark Philips NYU), and 5µg of pHR-PAG-GFP or pHR-PDZ-PAG-GFP. WT and PAG KO Jurkat T cells were transfected by incubating 2 x 10^6^ cells with 1mL of virus for 6 hours before being washed, cultured, and sorted for moderate GFP expression to match the PAG level with that of endogenous PAG in WT Jurkat T cells.

### Expressing WT-PAG-PAG and ΔPDZ-PAG-GFP in primary mouse T cells

Primary mouse CD8 T cells were transduced by retroviral transduction using a protocol adapted from as previously described (*68*). pMMLV retroviral constructs were generated by VectorBuilder with WT-PAG or ΔPDZ-PAG (4 C-terminal amino acids deleted) + unconjugated tdTomato (P2A linker) (VB221129-1260ayw, VB221205-1304djs). The virus was produced with Plat-E cells (Cell Biolabs RV-101) and transfected with 20µg of MMLV DNA, 43µL of Lipofectamine-3000, and 40µL of P3000 (Invitrogen L3000008). On day 2, the media was changed. On day 3, the virus was collected, filtered and immediately used for infection. A 24-well non-TC treated plate was coated with 5µg/mL anti-CD3 (BioLegend 100253) overnight at 4C or for 2h at 37C. 15µL of Retronectin (Takara Bio T100A) was added, and the plate was incubated for another 2h at 37C. At the end of the incubation, the Retronectin and anti-CD3 were aspirated, and 500µL of filtered virus was added to each well. The virus was spun onto the plate at 2000g at 32C for 1-2 hours.

Primary CD8 mouse T cells were isolated by negative selection (MojoSort for CD3 or CD8) and stimulated for 48 hours in a non-TC treated 24 well plates coated with 5µg/mL aCD3 overnight at 4C or for 2h at 37C. T cell media was supplemented with 2µg/mL aCD28 (BioLegend 102116). After 48h, T cells were harvested, washed, and resuspended in fresh T cell media supplemented with 6x IL-2 and 6x anti-CD28. After the 24-well plate was spun with the virus, 100µL of cells were added to each well and incubated with the virus for 48 hours. At 24h of incubation, 500µL of fresh T cell media was added to each well. After 48h, the cells were washed and sorted for tdTomato expression (BD Influx Cell Sorter with 405, 488, 561 and 638nm lasers). After sorting, cells were used immediately for adoptive cell transfer into mice or returned to culture at 37C with 5% CO2 in T cell media supplemented with 2.5µg/mL anti-CD3 and 1µg/mL anti-CD28 until use for in vitro assay. In the culture, new T cell media was added to the cells every 1-2 days to maintain optimal conditions.

### Synapse microscopy

All imaging was performed on ZEISS LSM 710 or ZEISS LSM 900 with a 20x or 63x objective. ImageJ (NIH, v1.54g) and Cell Profiler (Broad Institute, v4.2.1) were used for image analysis.

Raji B cells (ATCC) were pre-incubated with 2µg/mL SEE (Toxin Technology ET404) and co-cultured with Jurkat T cells. Cells were plated on poly-L-lysine-coated 35mm Mattek plates (Sigma-Alrich P8920; MATTEK P35G-1.5-14-C) and imaged live to observe synapse formation and localization of PAG-GFP and actin (LifeAct, AddGene 84385) within Jurkat T cells in the immune synapse.

To image the immune synapse parallel to the plane of imaging, Jurkat T cells were stimulated on Cellvis 8-chamber slides with #1.5 cover glass (Cellvis C8-1.5H-N) that had been coated with poly-L-lysine and pre-incubated with 5µg/mL anti-CD3 + 5µg/mL anti-CD28. Antibodies were washed off twice with complete RPMI, and 1 x 10^5^ PAG KO Jurkat T cells with WT-PAG-GFP or ΔPDZ-PAG GFP in 200µL complete RPMI were plated and placed in 37C 5% CO2 incubator for 5-40 min, at which point the cells were fixed by gently adding 200µL of warm 16% paraformaldehyde (Electron Microscopy Sciences 15700) in PEM buffer: 80mM kPIPES (Sigma-Aldrich P7643), 2mM MgCl_2_ (Sigma-Aldrich M8266), 5mM EGTA (Research Products International E14100), 2mM sucrose (Milipore 8590-OP) at pH 6.8 in PBS) (*69*). After 10-15 min of fixation at room temperature, cells were gently washed 3x with PBS by removing and replacing 50% of the volume each time. Cells were permeabilized for 5 min with 200µL of 0.25% Triton X-100 (Thermo Scientific Chemicals A16046.AE) in PBS and washed 3x with PBS, as above. Autoflorescence was quenched by adding 200µL freshly prepared NaBH4 (1mg/mL) in PBS to the cells for 7 min and then washed 3x with PBS, as above. Cells were blocked for 30 min with 200µL of 5% bovine serum albumin (BSA) (Sigma-Aldrich A9647) in PBS and then washed 3x with PBS, as above. Cells were stained with phalloidin-647 (AAT Bioquest 23127) in PBS for 60 min at room temperature in the dark. Cells were washed 3 x 5min with PBS, as above, and stored in PBS at 4C until imaging. Images were acquired on ZEISS LSM 900 microscope with 63x lens. The images were analyzed using Cell Profiler to mask cells based on actin staining and measure the area and perimeter of each masked cell.

### Co-immunoprecipitation and Western blot

2 x 10^6^ WT Jurkat T cells with stable expression of WT-PAG-GFP or ΔPDZ-PAG-GFP were washed twice with PBS and lysed for 20 min at 4C with gentle agitation in 300µL lysis buffer composed of 1:10 glycerol (Fisher Chemical G33-1), 1:20 1M Tris (Thermo Scientific Chemicals J62186.K2), 1:20 5M NaCl, 1:200 20% NP40, 1:250 0.5M EDTA (Invitrogen 15575020) + protease inhibitor (Roche 11-836-170-001). Lysates were then sonicated (Branson Digital Sonifier B450) at 25% power for 10 seconds and centrifuged (Eppendorf 5430R) at 12,000xg for 10 min at 4C. The supernatant was incubated with anti-GFP agarose beads (MBL D153-8) for 2h at 4C with gentle agitation. Beads were washed 4-5x and then incubated with Laemmli buffer Novex LC2676) at 95C for 3-5 min. Immunoprecipitate was run on a 4-20% Tris-glycine protein gel (Invitrogen XP04202BOX), and the gel was blotted onto a nitrocellulose membrane (Cytiva 10600002) with BioRad Trans-Blot Semi-Dry Transfer Cell (BIO-RAD 1703940) and Tris-Glycine Transfer Buffer (Thermo Scientific 28363) with 20% MeOH. The membrane was blocked with 5% BSA in 0.1% PBS-Tween 20 (Sigma-Aldrich P1379) for 20 min and incubated with rabbit anti-GFP (1:2000, Cell Signaling Technology 2956) and mouse anti-actin (1:5000, Invitrogen MA1-140) primary antibodies overnight at 4C. After 3x wash with 2.5% BSA in PBS-Tween, the membrane was incubated with goat anti-rabbit 680nm (1:10,000 Li-COR 926-68021) and goat anti-mouse 800nm (1:10,000, Li-COR 926-32210) secondary antibodies for 45 min at room temperature. The membrane was washed 3x, and bands were visualized and quantified on LI-COR Odyssey CLx Imager.

### Static adhesion assay

Static adhesion assay was modified from the protocol previously described (*70*). The assay was performed with primary mouse CD8 T cells that had been transduced with WT-PAG or ΔPDZ-PAG and sorted for tdTomato+ and then cultured with 2.5µg/mL aCD3 and 1µg/mL aCD28 antibodies. T cells were counted, washed, and resuspended in 1mL of staining solution. WT-PAG T cells were resuspended in 2µM CFSE (BioLegend 79898) in PBS, and ΔPDZ-PAG T cells were resuspended in 1µM Tagit-Violet (BioLegend 425101) for 10 min at room temperature in the dark. Cells were thoroughly washed and resuspended at 1 x 10^6^ cells/mL in T cell RPMI. 100µL of each cell type were combined and plated on Mattek plates that were pre-coated for 1h with 1:1000 mICAM1 (BioLegend 553004) in PBS. Plates were imaged at baseline or after one wash with T cell media that was performed after an incubation of 10, 20, or 30 min. Wash was performed gently to remove only cells without strong adherence to the plate. Imaging was performed with a 20x lens to capture two 6×4 tile regions per timepoint per sample. Images were analyzed using Cell Profiler to count the number of each cell type per 3×1 fields.

### Adhesion under shear stress

Adhesion under shear stress assay was modified from the protocol previously described (*50*). WT-CHO cells (Martin Sadowshi, NYU, New York) were cultured in 10mL F-12K media (ATCC 30-2004) supplemented with 10% FBS + 1% P/S in a 10cm plate at 37C with 5% CO2. One day before the experiment, CHO cells were trypsinized (Corning 25053CI), washed, counted, and resuspended at 2.5 x 10^6^ cells/mL in T cell media. 30µL of cells were slowly added to each chamber of the Ibidi flow chamber plate (Ibidi 80607). An additional 200µL of media was added, 50µL at a time, to the two reservoirs of each chamber. The flow plate was placed in a 10cm plate with 3mL of PBS to maintain humidity. The 10cm plate was incubated overnight at 37C with 5% CO2.

The adhesion assay was performed with primary mouse CD8 T cells that had been transduced with WT-PAG or ΔPDZ-PAG (described above) and sorted for tdTomato+ and then cultured with 2.5µg/mL aCD3 and 1µg/mL aCD28 antibodies. On the day of the experiment, T cells were counted, and for each chamber, 1 x 10^6^ of each cell type (WT-PAG and ΔPDZ-PAG) were washed and resuspended in 1 mL of staining solution. WT-PAG T cells were resuspended in 2µM CFSE in PBS, and ΔPDZ-PAG T cells were resuspended in 1µM Tagit-Violet for 10 min at room temperature in the dark. Cells were thoroughly washed and resuspended in 120µL of T cell media per 1 x 10^6^ cells. The two cell types were placed in the 37C incubator until use.

The microscope stage was heated to 37C and set to 5% CO2. Flow apparatus was constructed as described (*50*). Input tubing was flushed with water and then attached to a 60mL syringe filled with warm (37C) RPMI without FBS. Media was pushed through the tubing until it was filled with media. 60mL syringe was set in a syringe pump (New Era Instruments NE-300) to dispense 0.3mL/min. The Ibidi flow chamber was put on the microscope. 70µL of media was removed from the output reservoir, and 70mL of T cells were added to the input reservoir. This exchange was repeated three times to total 210uL. While the cells settle, the microscope focus was set on CHO cells using a 20x objective. 5-6 field positions were set and imaged for the baseline (0 min) timepoint with the 405 nm (Tagit-Violet) and 488 nm (CFSE) excitation lasers to capture both cell types. 10 min after cells were added, input and output tubing were carefully attached to avoid air bubbles, and the syringe pump was started to allow flow through the chamber at 0.3mL/min. After 5 min of flow, the syringe pump was stopped, and the exact positions were imaged again. Images were analyzed using a Cell Profiler to count the number of each cell type per field.

### Degranulation Assay

The protocol was adapted from the previously described (*71*). Primary mouse CD8 T cells that were transduced with WT-PAG or ΔPDZ-PAG (as described above) and sorted for tdTomato expression were cultured with CFSE-stained MC38-OVA cells (Jun Wang, NYU) at a 1:1 ratio at 37C with 5% CO2 in a U-bottom 96-well plate in 200µL of T cell media with 10µL of anti-mouse CD107a-BV421 (BioLegend 121617). After 1 hour, 1:500 of protein transport inhibitor cocktail (Invitrogen 00-4980-93) was added, and cells were cultured for another 3 hours. Cells were transferred to a FACS tube, washed twice with PBS, and stained with Zombie NIR (BioLegend 423106). Cells were analyzed with BD LSR-II flow cytometer (BD Biosciences, UV, 405, 488, 532, 594 and 633nm lasers) and FlowJo software (BD Biosciences, v10.9).

### In vitro Cytotoxicity assay

Raji B cells were washed and stained with Tagit-violet and live cell dye Calcein (BioLegend 425201) for 15 min. Cells were counted, washed 2x, and resuspended at 1 x 10^6^ cells/mL in T cell media supplemented with 100ng/mL SEE and 100ng/mL SEB (Toxin Technology BT202) for 30 min before adding prepared T cells. To prepare T cells, primary mouse CD8 T cells were transduced with WT-PAG or ΔPDZ-PAG (as described above) and sorted for tdTomato expression. They were cultured overnight with 2.5µg/mL anti-CD3 and 1µg/mL anti-CD28 in T cell media. T cells were counted, washed, and resuspended in CellTrace FarRed (Invitrogen C34572) 1:1000 in PBS for 15 min. After staining, T cells were washed 2x and resuspended at 1 x 10^6^ cells/mL in T cell media. T cells were added to Raji cells at 1:1 ratio and cultured on poly-L-lysine coated Cellvis 8-chamber slides with #1.5 cover glass for 24h at 37C with 5% CO2.

After 24h, plates were imaged on a ZEISS LSM 900 microscope using the 20x lens to capture 9-11 frames of Tagit-Violet (Raji), Calcein (live Raji), and CellTrace Far Red (T cells). Images were analyzed with ImageJ to measure the total signal in the Violet and Calcein channels. The calcein signal was normalized to the Violet signal in each frame to normalize the number of Raji cells in the frame. The normalized calcein signal was compared to the normalized calcein signal from Raji cells cultured alone without T cells to determine the relative decrease in calcein signal and thus the percent of Raji cell death when cultured with CD8 T cells.

### Type IV Hypersensitivity Model

Animal studies were approved by the Columbia University Institutional Animal Care and Use Committee. As described above, MMLV-WT-PAG and MMLV-ΔPDZ-PAG viruses were produced and transduced into preactivated CD3-isolated mouse splenocytes. After 48h incubation with viruses, cells were harvested, tdTomato expression was assessed by flow cytometry, and an equivalent number of cells was transferred into PAG KO mice by tail vein injection. One day following adoptive cell transfer, the abdomens of the recipient mice were shaved and treated with 100µL of 0.5% DNFB (Sigma-Aldrich D1529) in 4:1 acetone (LabChem LC104202): olive oil (Sigma-Alrich O1514). Shaving and application was performed under isoflurane anesthesia to facilitate absorption. Mice were checked daily, and five days later, they were challenged with the application of 30µL of 0.3% DNFB to the back of their left ear. The application was performed under anesthesia to facilitate absorption. At 48h after the challenge, mice were euthanized, and the thickness of both ears was measured with a digital micrometer (Beslands B07KC7YNR8) three times each. The tissue was flushed with PBS by transcardiac perfusion. Ears and abdominal skin were fixed in 10% formalin (Epredia 5725) for 24-48 hours and then transferred to 70% histology-grade ethanol (Fisher Chemical A405P-4) until further processing and assessment by histology. Spleen, inguinal LNs (primary site draining LN), and cervical LNs (challenge site draining LN) were removed, weighed (Mettler Toledo XSR205), and stored in 10% FBS-PBS on ice until processing and assessment by flow cytometry.

### Tumor Model

Animal studies were approved by the Columbia University Institutional Animal Care and Use Committee. 6–8-week-old male C57BL/6J mice with CD45.1 (JAXBoy— Ptprc^K302E^; Jackson Labs Strain # 033076) were implanted intradermally with 2.5-3 x 10^6^ MC38-OVA cells in the right hind flank. Mice were monitored daily, and tumor measurements were taken by digital caliper (NEIKO 01417A) every 1-2 days beginning 3-4 days after implantation. Tumor volume was approximated using the following formula: V = ½ length ∈ width^2^. On day 14 or 17, mice received adoptive transfer of CD8 T cells isolated from PAG-KO OT-1+ CD45.2 mice that were transduced with MMLV-WT-PAG or ΔPDZ-PAG, as described above, and sorted for tdTomato expression. Mice continued daily monitoring and tumor measurements until they were euthanized 9-12 days after adoptive cell transfer. The tissue was flushed with PBS by transcardiac perfusion. Inguinal LNs, spleen, and tumor were removed, weighed, and stored in 10% FBS-PBS on ice until processing and assessment by flow cytometry. A sample of tumor tissue was fixed in 10% formalin for 24-48 hours and transferred to 70% histology-grade ethanol until further processing and assessment by histology.

### Histology

Tissues were processed and embedded in paraffin blocks. Slides were either stained with H & E (MPSR CUMC, New York) or IHC. For IHC, 5µm tissue sections were deparaffinized with xylene (Sigma-Aldrich 534056) and rehydrated with a gradient of ethanol solutions. Slides were immersed in IHC Antigen Retrieval Solution, low pH (Invitrogen 00-4955-58), and microwaved for two cycles of 100% power for 2 min followed by 20% power for 15 min. Slides were washed 3x in TBS (Boston BioProducts BM-300), blocked in 1% BSA in TBS for 45 min, and washed 2 x 5 min. Slides were incubated with 1:400 primary antibody in TBS for 90 min at room temperature or overnight at 4C. Primary antibodies used were rabbit anti-mouse CD3 (Abcam ab16669) and rabbit anti-RFP (Rockland 600-401-379). Slides were washed 3 x 5 min and stained with 1:1 HRP-conjugated goat anti-rabbit IgG (Abcam ab214880) in 1% BSA-TBS for 2 hours at room temperature. Slides were washed 3 x 5 min and stained for 10 min with 1:50 DAB Chromogen in DAB substrate (Abcam ab64238). Slides were rinsed 4x, stained with hematoxylin (Sigma-Aldrich GHS316), rewashed 4x, and mounted with aqueous mounting medium for IHC (Abcam ab64230). Slides were sealed and scanned with an Aperio AT2 Slide Scanner microscope (Leica Biosystems).

### Flow Cytometry

LN and spleen tissue was dissociated through a 70-micron filter using the back of a 5mL syringe plunger and rinsing with 10% FBS-PBS. Dissociated cells were washed once. The spleen pellet was resuspended in 1-2mL of ACK lysis buffer (KD Medical RGF3015), incubated for 5 min, and washed again with 10% FBS-PBS. Tumor tissue was chopped into small pieces in 3-5mL of digestion buffer: 1mg/mL collagenase (Sigma-Adrich C0130), 20 µg/mL DNase I (STEMCELL Technologies 7469), and 10% FBS in RPMI or PBS. The tissue was incubated in a digestion buffer at 37C for 30 min with periodic mixing. Digested tissue was vortexed and dissociated through a 40-micron filter (Corning 352340) using the back of a 5mL syringe plunger and the large end of a p1000 pipet tip. Dissociated cells were washed once and filtered again through a 40-micron filter. Cells were washed and resuspended in 5mL of 40% Percoll (Cytiva 17089102) in PBS in a 15mL tube and spun at 1000g for 10 min at room temperature with acceleration and brake set to 1 (Eppendorf 5810R). The supernatant was discarded, and the pellet was resuspended in 10% FBS-PBS. Cells were counted, and up 5 x 10^6^ cells were taken for staining.

For hypersensitivity experiments, samples were transferred into 5mL FACS tubes and washed with PBS. They were first stained with 1:250 Zombie UV (BioLegend 423108) and 1:50 TruStain FcX anti-mouse (BioLegend, 101320) in 50mL PBS for 10 min at room temperature. Then cells were incubated in the dark for 20 minutes with 50µL of the antibody mixture: 1:50 each of anti-CD3-AF488 (BioLegend 100210), anti-CD4-BV510 (BioLegend 100559), anti-CD8-PerCP/Cy5.5 (BioLegend 140418), anti-CD44-BV421 (BioLegend 103040), anti-CD62L-BV711 (BioLegend 104445), anti-CD69-APC (BioLegend 104514), and anti-PD1-PE/Cy7 (BioLegend 109110). Cells were washed twice and resuspended in 1:4 fixation buffer (BioLegend 420801) in PBS for 10 min at room temperature. Samples were washed twice and stored at 4C until analyzed by flow cytometry using BD LSR-II. Results were analyzed using FlowJo.

For tumor experiments, samples were transferred into a V-bottom 96-well plate and washed once with PBS. They were first stained with 50µL of 1:200 Zombie NIR in PBS at room temperature for 30 min. Cells were washed with 200µL of staining buffer: 10g/L BSA, 1:100 sodium azide (Bioworld 41920044-1), and 1:200 EDTA (stock at 0.5M) in PBS. Next, cells were incubated with 50µL of 1:50 TruStain FcX anti-mouse (BioLegend, 101320) for 10 min at room temperature. Cells were washed with 200µL of staining buffer. Samples were resuspended in 100µL of antibody mixture: per sample, staining buffer with 2.5µL SuperBright (Invitrogen SB-4401-75), 2.5µL CellBlox (Invitrogen B001T06F01), 0.02µL anti-CD4-BUV615 (BD Biosciences 613006), 1.25µL anti-LAG3-BUV805 (BD Biosciences 748540), 0.05µL anti-PD1-BV480 (BD Biosciences 568612), 2.5µL anti-CD25-BV510 (BD Biosciences 563037), 0.15µL anti-CD8-BV570 (BioLegend 100740), 0.3µL anti-CXCR3-BV750 (BD Biosciences 747298), 1.25µL anti-CX3CR1-BB515 (BD Biosciences 567809), 1.25µL anti-CD45.2-PerCP/Cy5.5 (BioLegend 109827), 0.02µL anti-MHCII-RY586 (BD Biosciences 568546), and 0.15µL anti-CD45.1-PE/Cy7 (BioLegend 110729). Samples were incubated with the antibody mixture for 40 minutes at room temperature in the dark. Cells were washed once with 150uL of staining buffer and twice more with 250µL of staining buffer. Cells were resuspended in 100µL of 1:4 Fixation/Permeabilization concentrate (Invitrogen 00-5123-43) in Fixation/Permeabilization diluent (Invitrogen 00-5223-56) and incubated for 45 min at room temperature in the dark. Samples were washed 2x with PBS and kept at 4C overnight. The next day, samples were spun down and washed with 250uL of 1:10 permeabilization buffer concentrate (Invitrogen 00-8333-56) in water. Samples were incubated for 45 min at room temperature in the dark with 100µL of intracellular antibody mixture: per sample, 1x permeabilization buffer with 2.5µL SuperBright, 2.5µL CellBlox, 0.2µL anti-Ki67-RB780 (BD Biosciences 568761), 0.05µL anti-GranzymeB-APC (BioLegend 372203), and 0.3µL anti-CD3-SparkRed718 (BioLegend 100282) for 45 min at room temperature in the dark. The plate was washed once in 150µL of permeabilization buffer and once in 250uL of permeabilization buffer. Cells were resuspended in 200uL of PBS and transferred to 5mL FACS tubes for data collection with a 5-laser Aurora spectral flow cytometer (CYTEK). Results were analyzed with FCS Express (De Novo Software, v7).

### Statistics

All results represent a minimum of two independent experiments. Bar graphs and tumor growth curves show mean and standard deviation. Box and whisker plots show a box from the 25^th^ to 75^th^ percentile with a line at the median and whiskers from 5^th^ to 95^th^ percentile. Unless otherwise stated in the figure legend, all analyses were chi-square (Fig 1), Mann-Whitney t-tests for samples with n < 15, or unpaired t-tests for samples with n ≥ 15. Multiple comparisons were performed by ordinary one-way ANOVA. All statistical analyses were performed using Prism (GraphPad, v10). Principal components analysis (PCA) was performed using GraphPad Prism 10 with MFI values from flow cytometry results, normalized between experiments, and set to range from 0-100 for each parameter. Uniform Manifold Approximation and Projection (UMAP) analysis was performed using FlowJo with FlowSOM supervised clustering of 13 parameters into 9 clusters.

## Supplementary Materials

Fig. S1. Validation of PAG expression.

Fig. S2. DNFB hypersensitivity gross measurements. Fig. S3. MC38-OVA tumor and in vitro stimulation.

## Acknowledgments

We would like to thank Mark Philips (New York University, New York, USA) for providing lentiviral vectors, Martin Sadowski (New York University, New York, USA) for providing cell lines, and Alexander Tarakhovsky (The Rockefeller Institute, New York, USA) for providing PAG-KO mouse strains, all of which were critical to this study. Several figures were created using BioRender.

## Author contributions

Conceptualization: EKM, MS, AM

Formal Analysis: EKM

Funding Acquisition: AM, EKM, EMM

Investigation: EKM, HX, CT, XX

Methodology: EKM, MS, HX, CT, MP, MJS, SB, SL

Project administration: AM, EKM

Resources: AM, EMM, RW

Supervision: AM, MS, MP, SB, SL, EMM, RW

Visualization: EKM

Writing—original draft: EKM, AM

Writing—review and editing: EKM, AM, MS, EMM, MJS, MP

## Data and materials availability

All data is available in the main text or the supplementary materials. All primary data will be made available upon request to the corresponding author.

