## Supplemental Figures for "PAG orchestrates T cell immune synapse function by binding to actin"

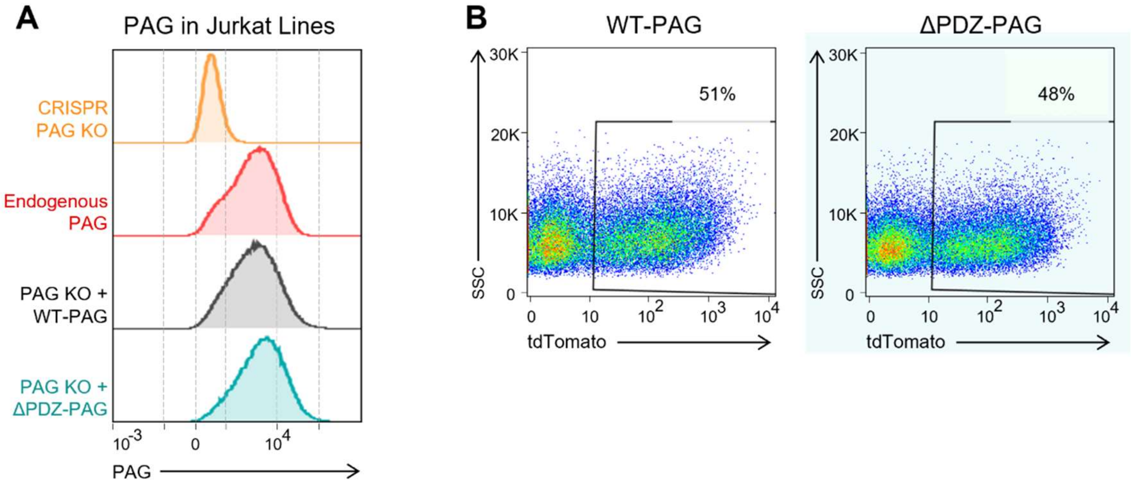


**Fig. S1. Validation of PAG expression. (A)** Jurkat T cells were depleted of PAG by CRISPR and then stably transduced to re-express WT-PAG-GFP or ΔPDZ-PAG GFP. Cells were fixed, permeabilized, and stained intracellularly for PAG to compare PAG levels between PAG KO, wildtype (endogenous), PAG KO + WT-PAG-GFP rescue, and PAG-KO + ΔPDZ-PAG-GFP rescue. PAG levels were measured by flow cytometry after gating on Live cells. **(B)** Primary mouse T cells from PAG-KO mice were transduced with WT-PAG + tdTomato or ΔPDZ-PAG + tdTomato, and tdTomato expression was evaluated by flow cytometry. Plots show tdTomato expression before the cells were sorted according to the gates shown.

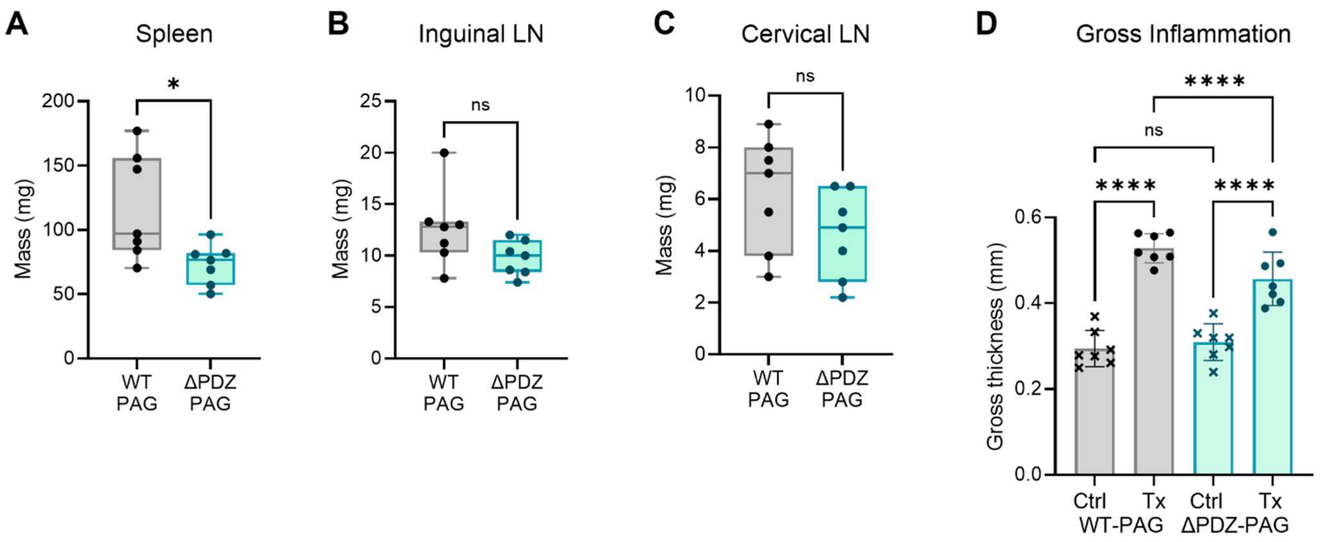


**Fig. S2. DNFB Hypersensitivity Gross Measurements.** Mice underwent a DNFB hypersensitivity experiment. N = 6-7 mice per group across 3 independent experiments. After euthanasia, the mass was taken for the **(A)** spleen, **(B)** inguinal LNs, and **(C)** cervical LNs. Statistics by Mann-Whitney t-test. **(D)** At the endpoint of the DNFB hypersensitivity experiment, the thickness of each ear was averaged from three measurements by digital caliper. Statistics by ANOVA with Sidak's test for multiple comparisons. **ns** = P > 0.05, ∗ = P ≤ 0.05, ∗∗∗∗ = P ≤ 0.0001.

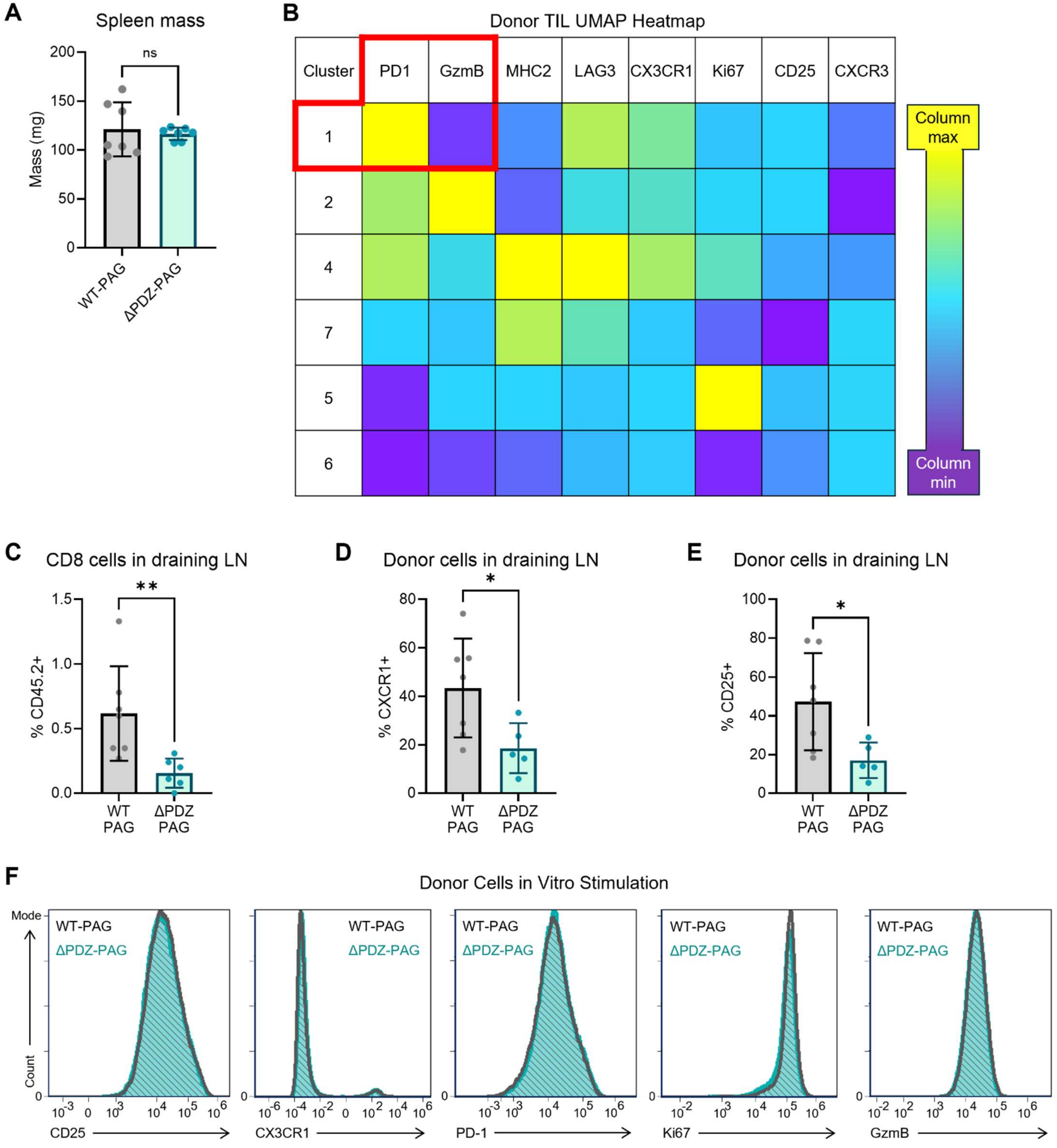


**Fig. S3. MC38-OVA tumor LN and in vitro stimulation. (A-E)** Mice underwent experiment with MC38-OVA tumors and adoptive transfer of OT-1 T cells expressing

WT-PAG or ΔPDZ-PAG. N = 7 mice per group across 3 independent experiments. **(A)** After euthanasia, the mass was taken for the spleen. **(B)** Unsupervised cluster analysis and UMAP visualization was performed on flow cytometry results from donor TILs. Protein expression relative to all other clusters is shown, where yellow is the cluster with the highest level of the column's protein, and purple is the lowest expression level. The six most populated clusters are shown. **(C-E)** The cells in the tumor-draining LN (inguinal) were assessed by flow cytometry. Statistics by Mann-Whitney t-test. **(C)** Lymphocytes were gated on CD3+CD8+ cells. The percent of these cells with CD45.2 (donor cells) is shown. Expression of **(D)** CX3CR1 and **(E)** CD25 within the population of donor CD8 T cells isolated from the tumor-draining LN. **(F)** After sorting for WT-PAG or ΔPDZ-PAG expression based on tdTomato fluorescence, donor OT-1 T cells were cultured in vitro with stimulation by soluble anti-CD3 and anti-CD28 antibodies. The cells were assessed by flow cytometry for expression of the following proteins: CD25, CX3CR1, PD-1, Ki67, and Granzyme B. Representative results are shown from N = 3 independent experiments. **ns** = $P > 0.05$, ∗ = $P \leq 0.05$, ∗∗ = $P \leq 0.01$.
